# Mutational divergence over years in local populations of the selfing nematode *Caenorhabditis elegans*

**DOI:** 10.1101/2025.11.19.689209

**Authors:** Xu Wei, Aurélien Richaud, Robyn E. Tanny, Erik C. Andersen, Marie-Anne Félix

## Abstract

Laboratory mutation accumulation experiments allow the assessment of spontaneous mutation rates and patterns with minimal selection. Here, we aimed to follow the accumulation and fate of mutations in natural populations, in a spatial context. The nematode *Caenorhabditis elegans* is particularly suited for such endeavor, as it reproduces almost exclusively by selfing. We analyzed the evolution of clonal *C. elegans* genotypes along a 300-m long stream bank in the Santeuil wood (France), based on short-read whole-genome sequencing of individuals collected between 2009 and 2022. We followed along years two of these quasi-clones composed of individuals only differing by recent mutations. Recombination was scarce. A temporal signal was detected: strains from earlier years were found close to inner nodes of the tree, while recent ones were found on outer tips. This signal allowed us to estimate a substitution rate of 4 to 5×10^-8^ mutations per base pair per year, which can be used to calibrate divergence times among and within species. Mutation densities were higher on the X chromosome, on chromosome arms, and in non-exonic regions. We detected a high transition-to-transversion ratio, not observed in *C. elegans* laboratory mutation accumulation lines. Based on the spontaneous mutation rate per generation in laboratory lines of intergenic regions under minimal purifying selection, we estimated that *C. elegans* locally undergoes around 25 effective generations per year. Finally, using these recent mutations, we detected a spatio-temporal pattern within the field site, indicating limited dispersal at the scale of 100 meters within 10 years.

## Introduction

Genetic studies of natural populations include evolutionary processes across wide timescales. In a typical population, the genealogy of genotypes integrates numerous recombination events on chromosome fragments, each carrying polymorphisms that arose by mutation across millions of years. Often, the derived allele of these polymorphisms cannot be inferred. At the other end of the spectrum, at very short timescales, isogenized populations in the laboratory can be cultured at minimal population size for several generations of mutation accumulation, allowing to assess rate, patterns and phenotypic effects of spontaneous mutation with minimal selection (Lynch 1985; Keightley 1994; Keightley and Caballero 1997; Vassilieva and Lynch 1999; Denver *et al*. 2009; Konrad *et al*. 2019; Saxena *et al*. 2019). Immediately above this laboratory timescale, we aimed here to study the mutational process in natural populations that are subject to selection, in their environment. Some organisms naturally provide isogenic populations where mutation can be easily studied in a given genomic background. However, most such species do recombine at a non-negligible rate. For example, extensive genomic surveillance has documented the accumulation of spontaneous mutations in viruses, yet re-assortment and recombination events (Holmes *et al*. 2005; Shao *et al*. 2017; Roemer *et al*. 2023) complicate the inference of mutational trajectories. The crucifer *Arabidopsis thaliana* underwent a narrow initial population bottleneck during a colonization event of the American continent a few centuries ago. Using whole-genome sequencing of museum samples, (Exposito-Alonso *et al*. 2018) leveraged this bottleneck to analyze the advent and spread of new mutations in a clonal natural lineage with rare recombination.

Studying timescales over many generations is facilitated with organisms displaying short generation times. The nematode *Caenorhabditis elegans* provides an excellent animal system where the appearance and fate of mutations can be directly studied in the wild. Given its fast lifecycle, *de novo* mutation can be detected and studied on the timescale of a few years rather than centuries. *C. elegans* reproduces mostly through selfing XX hermaphrodites (modified females that produce sperm early in adulthood) and rare X0 males arising by spontaneous non-disjunction of the X chromosome or as progeny of a cross. Outcrossing is rare in most natural populations where it was evaluated, and distinct independent clones co-exist in the same location (BarriÈre and FÉlix 2005; Haber *et al*. 2005; Sivasundar and Hey 2005; BarriÈre and FÉlix 2007; Richaud *et al*. 2018; Crombie *et al*. 2019; Crombie *et al*. 2022). Based on the comparison between short-term and long-term measures of outcrossing, the cross progeny of distinct genotypes even appear to be counterselected (BarriÈre and FÉlix 2007). This rare effective recombination in selfing *Caenorhabditis* nematodes could be explained by outbreeding depression (Dolgin *et al*. 2007; Gimond *et al*. 2013) and gene drive effects (Seidel *et al*. 2008; Ben-David *et al*. 2017; Noble *et al*. 2021). At the species level, each group of highly similar genotypes, defined by a threshold of genomic similarity (Crombie *et al*. 2024), is called an ‘isotype’ (Andersen *et al*. 2012). Several of these isotypes co-exist locally (BarriÈre and FÉlix 2005; Haber *et al*. 2005; Sivasundar and Hey 2005; BarriÈre and FÉlix 2007; Richaud *et al*. 2018; Crombie *et al*. 2022). Here we focus on local polymorphisms *within* an isotype, which have not been studied so far.

Several motivations led us to study the accumulation of mutations in a natural population. First, dating the divergence between nematode species is a challenge, because nematodes have notoriously long branches on animal phylogenies and are poorly fossilized. For example, some *Caenorhabditis* species are as distant in terms of protein sequence evolution than fish and human (Kiontke *et al*. 2004), yet only represent a small fraction of nematode diversity. Methods that estimate *Caenorhabditis* species divergence dates require an estimate of the substitution rate per year (Cutter 2008; Fusca *et al*. 2025; Picao-Osorio *et al*. 2025), which we aimed to provide here in a natural context for *C. elegans*. Second, by calibrating the nucleotide substitution rate with a mutation rate per generation estimated in the laboratory could allow us to estimate the number of effective generations per year for the species. The lifecycle of *C. elegans* can be completed within three days in favorable conditions, but in real life development can be arrested at various points, including in a modified third larval stage called the dauer, commonly found in natural populations (BarriÈre and FÉlix 2005; BarriÈre and FÉlix 2007; FÉlix and Duveau 2012; Richaud *et al*. 2018). This raises a question about *C. elegans*’ generation time in the wild and the effective number of generations the species undergoes per year. Third, following the accumulation of mutations over time and space could allow us to track movement of sublineages of these microscopic animals. Aims of the present study were thus to estimate the rate of mutations and the number of generations per year in a natural population, and to track mutations in a spatial context.

*C. elegans* thrives in patchy bacteria-rich natural habitats provided by decomposing plant substrates, such as stems of herbaceous plants, fruits and flowers (FÉlix and Braendle 2010; FÉlix and Duveau 2012; FrÉzal and FÉlix 2015; Dirksen *et al*. 2016; Samuel *et al*. 2016; Schulenburg and FÉlix 2017; Crombie *et al*. 2022). The species is found throughout the world, mostly in temperate regions, and at a higher elevation in tropical regions (Andersen *et al*. 2012; Crombie *et al*. 2019). To investigate within-isotype polymorphisms, we chose to focus on the Santeuil wood in France, a site for *Caenorhabditis* collection since 2009 (FÉlix and Duveau 2012; Richaud *et al*. 2018). The field site consists in a 300-meter-long North-South transect bordered by a small stream on the East side (the Viosne, running North to South), a grass field on the North end, a wood with little undercover of herbaceous plants on the West side and a stream tributary on the South end (Figure 1a,b). We collected samples of decomposing plant material, mostly stems of herbaceous plants such as *Heracleum, Symphytum, Tussilago*, as well as some invertebrates that presumably carry *C. elegans* between bacterial-rich food sources (Figure 1c) (BarriÈre and FÉlix 2005; Caswell-Chen *et al*. 2005; FÉlix and Duveau 2012; Petersen *et al*. 2015; Schulenburg and FÉlix 2017). Demographic and developmental stage compositions was the focus of the first study (FÉlix and Duveau 2012). In the second study, three local isotypes named HS1, HS2, and HS3 (HS# for Haplotype Santeuil) were found using partial genome sequencing of animals collected from 2009 to 2014 (Richaud *et al*. 2018). These isotypes are not merely recombinant of each other and hardly ever crossed.

**Figure 1.**
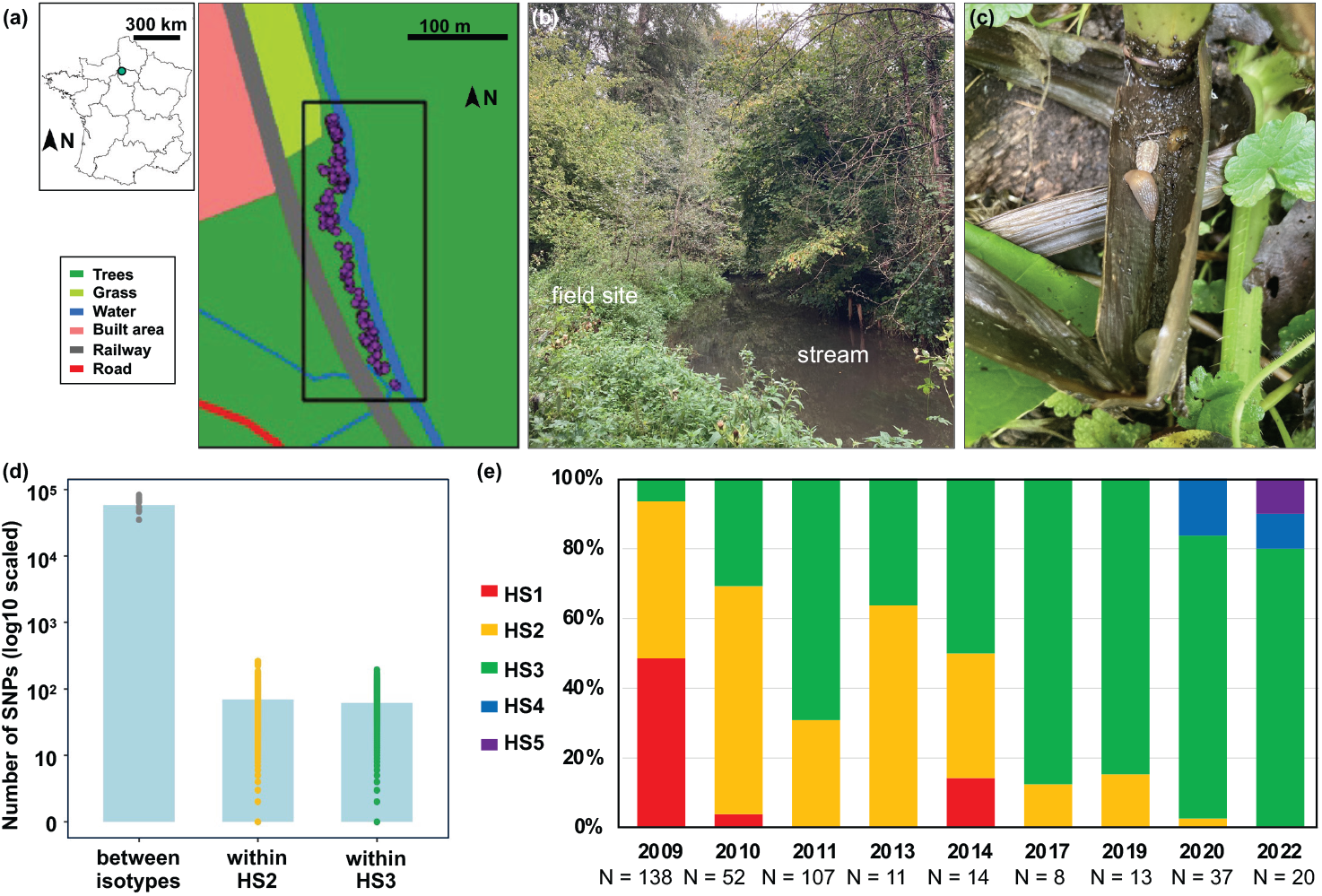
The Santeuil collection site and isotype proportions over years. (a) In the top left corner, location of Santeuil in mainland France, indicated by a green dot. The detailed map shows the sampling transect with the surrounding landscape structure. The land cover refers to ESRI (2021) and OpenStreetMap (2021). The purple dots indicate samples. (b) Picture of part of the sampling site along the stream, seen from the South. (c) Picture of relevant substrates (e.g., rotting stems and invertebrate animals, here slug and isopod) sampled in the field. (d) Numbers of SNPs between isotypes and within isotypes. (e) Population composition of *Caenorhabditis elegans* in the sampling line across years). The sample sizes (N) refer to the 2b-RAD dataset in (Richaud *et al*. 2018) (samples collected from 2009 to 2014) and whole-genome sequencing from CaeNDR (Crombie *et al*. 2024) (2017-2020) and the laboratory (2022).

The local isotype set in Santeuil provides an exemplary system to study the mutational landscape and spatial structuring of natural populations. Based on short-read sequencing of the whole genomes of strains collected between 2009 and 2022, we aimed to address the following questions: What are the mutation rate and patterns in natural populations of *C. elegans*? How many effective generations per year does *C. elegans* reproduce in the field? What are the spatiotemporal distribution patterns of *C. elegans* at this local scale?

## Materials and Methods

### Field collection

*Caenorhabditis elegans* were collected along the Viosne stream in the Santeuil wood as previously described (FÉlix and Duveau 2012; Richaud *et al*. 2018). This stream border includes herbaceous plants of humid areas, such as *Symphytum officinale* (comfrey, Boraginaceae) or *Tussilago farfara* (coltsfoot, Asteraceae), as well as *Heracleum sphondylium* (common hogweed, Apiaceae) and one or several unidentified taller species in the Asteraceae family. We also collected potential invertebrate vectors such as slugs, snails, isopods, insects and annelids. We recorded at the time of collection the position of each sample along the North-South transect using GPS (on a specific hand-held device in earlier years and a smartphone in later years). In addition, we used manual notes to precisely locate samples collected at distances of less than 10 meters (FÉlix and Duveau 2012; Richaud *et al*. 2018). We collected new strains for this work in 2017, 2019, 2020, and 2022. The 2020 collection focused on collecting stems and their associated invertebrates. A list of analyzed strains is provided in Table S1.

Strains were established by selfing for 3-5 generations of single wild-born individuals and assigned a unique identifier (e.g. JU1793). A single heterozygote between HS2 and HS3 had been found in (Richaud et al. 2018), which was not used here. Given freezing and transfer between laboratories, the strains may have undergone around 10-15 generations of selfing by the time of sequencing; note that we removed the terminal tree branches in estimating the mutation rate, and thus the contribution of these generations - see below. The HS1, HS2, and HS3 isotypes (Richaud et al. 2018) correspond to isotypes JU1934, JU1793, and JU2600 in CaeNDR (the *Caenorhabditis* Natural Diversity Resource; (Crombie *et al*. 2024), respectively. We sequenced and analyzed HS2 and HS3 strains from field collection spanning 2009 to 2022, usually a single strain per positive sample. Many more lines were frozen in (FÉlix and Duveau 2012; Richaud *et al*. 2018) and many genotyped by 2b-RAD sequencing in (Richaud *et al*. 2018).

### Genomic DNA sequencing

For many strains, DNA sequence alignments were extracted from the CaeNDR website version 20231213 (Crombie *et al*. 2024). For the strains from 2022, nematode culture, harvesting, and DNA extraction were performed as in (Richaud *et al*. 2018). Bleached animals were cultured at 20°C on large OP50-seeded NGM plates. Once the plates approached starvation (no visible *E. coli* OP50), *C. elegans* were harvested with M9, pelleted (5 min, 5,000 rpm), and washed once more. Genomic DNA was sequenced with 150 bp paired-end reads using Illumina sequencing by Eurofins Genomics. The raw sequences of these strains are available at NCBI Sequence Read Archive under accession PRJNA1280162.

### Read mapping and calling of single-nucleotide polymorphisms

This study focused on single-nucleotide polymorphism (SNP) in the nuclear genome. The *C. elegans* haploid nuclear genome size is 100 Mb, distributed on six chromosomes (Consortium 1998). The pipeline is summarized in Figure S1.

New Illumina reads were trimmed and aligned to the *C. elegans* reference genome WS283 (Sternberg *et al*. 2024) by fastp v0.20.0 (Chen *et al*. 2018) and BWA v0.7.17 (Li and Durbin 2009). The sequence alignments were then sorted and marked by sambamba v1.0.0 (Tarasov *et al*. 2015) and picard v2.26.2 (Institute 2021). Variant calling was performed using the sequence alignments (.bam files) of all strains identified at CaeNDR (version 20231213) as belonging to the isotypes JU1934 (HS1), JU1793 (HS2), JU2600 (HS3) and JU4161 (HS4) using GATK v4.2.2.0 (McKenna *et al*. 2010). Parametrization of read processing and variant calling were based on those used at CaeNDR (Crombie *et al*. 2024) (version 20231213; https://github.com/AndersenLab). The code is available as File S1. Briefly, GVCF files were first generated using GenotypeGVCFs tool with default parameters. The resulting VCF was processed with het_polarization and vcffixup in vcflib (Garrison *et al*. 2022). Following filtering (site-level: QUAL < 30, FS > 100, QD < 20, SOR > 5; genotype-level: DP < 5), a last filter was applied to retain for subsequent analysis only biallelic homozygous sites with a genotype depth (DP) ≥ 6.

We found in the 2022 samples a new Santeuil genotype HS5, represented for example by strain JU4370. We compared it to all isotypes at CaeNDR and found it to be novel, above the threshold of 99.97% of the overall SNP set. This isotype is now included in CaeNDR (version 20250625). For Figure 1, we evaluated the proportion of the different isotypes for each year of sampling using i) 2b-RAD data from (Richaud *et al*. 2018) covering years from 2009 to 2014 (as they included many more individuals) and ii) whole-genome sequences for the new field collections in 2017, 2019, 2020, and 2022. The proportions of isotypes are provided in Table S2.

Mutation rates and patterns were analyzed separately for isotypes HS2 and HS3. All strains either found in the Santeuil wood (thereafter “local”) or other locations outside the wood (“external”) in each isotype (i.e., HS2 and HS3) were employed for accurately calling variants. We trimmed the genome (called *trimmed genome*) to eliminate highly repetitive (masked regions in WormBase, (Sternberg *et al*. 2024) and hyper-divergent regions relative to the N2 reference strain (Crombie *et al*. 2024), as well as regions with read depths less than six to avoid miscalling. To control for potential confounding effects of novel mutations arising in the laboratory, we removed mutations in the heterozygous state. This conservative filter may exclude true, low-frequency heterozygous variants from the natural population, potentially leading to underestimates of genetic diversity and reduced power to detect recent or weak selection. However, because a key aim of this study was to estimate population genetic parameters such as mutation rate and effective generation number, we prioritized the use of variants originating from natural population history over those that were likely to reflect laboratory mutations or mapping errors. SNP sets are provided in Tables S3 and S4 for HS2 and HS3, respectively.

### Haplotype network construction and recombination

Haplotype networks were built using all known strains belonging to HS2 or HS3 by SplitsTree v6.2.1-beta (Huson and Bryant 2006). Three recombinants were identified within HS2 (Figure S3). For each recombinant, the recombination breakpoints on the chromosomes were inferred based on the distribution of SNPs.

### Mutation spectra

Based on their locations in exon, intron, non-coding gene, or intergenic regions, SNPs of local strains were categorized into four groups. Their distribution was visualized on the trimmed genome. Trimmed genomes and category sizes are provided in Table S5. SNP density was calculated across the trimmed genome in the following contexts: (1) per chromosome; (2) in tips, arms, and core regions predicted by (Rockman and Kruglyak 2009); and (3) in exon, intron, non-coding gene, and intergenic regions. SNP density was calculated by dividing the number of detected SNP loci within a region of interest (e.g., exon regions or the entire genome) by the total base pair content of that region. We identified nine and 27 dinucleotide variants in HS2 and HS3, respectively. For the purpose of these density calculations, dinucleotide variants were lumped as a single site. Subsequently, Chi-squared goodness-of-fit tests were employed to assess whether SNPs were evenly distributed among the categories within each context. With dinucleotide variants treated as two separate sites, we calculated the densities of all transition and transversion types and the *Ts/Tv* ratio across exonic, intronic, non-coding gene, and intergenic regions.

### SNP annotation

Annotation with respect to protein sequence was performed by csq function (Danecek and McCarthy 2017) of bcftools v1.13 (Danecek *et al*. 2021). The SNPs located in coding regions (CDS) were categorized into non-synonymous and synonymous mutations. A Chi-squared test was used to assess whether the distribution of non-synonymous and synonymous mutations differed significantly between wild strains from Santeuil and mutation accumulation lines maintained in the laboratory (Denver *et al*. 2009; Denver *et al*. 2012; Konrad *et al*. 2019).

### Phylogenetic analyses and estimation of substitutions and effective generations per year

To estimate local substitution rates in the *C. elegans* population from Santeuil, we implemented a Skygrid coalescent model (Gill *et al*. 2013) in BEAST v1.10.4 (Drummond and Rambaut 2007). For each isotype (HS2 and HS3), all SNPs from local non-recombinant strains were partitioned into four categories: exonic, intronic, non-coding gene-related, and intergenic regions. A total of 10,000 invariant sites were randomly generated and evenly distributed among these four categories to meet the base frequency requirements of the substitution model. Tip dates were calibrated based on sampling time. Each partition was assigned an uncorrelated relaxed clock model (lognormal distribution) and a GTR+G+I substitution model, allowing rates to vary across branches. To minimize the influence of singleton mutations originating from recent laboratory mutations on substitution rate estimates, we focused on the mean evolutionary rate along internal branches. The MCMC analysis was run for 100 million generations, sampling 10,000 steps. A Maximum Clade Credibility (MCC) tree was summarized using the TreeAnnotator module in BEAST for visualization.

Referring to the substitution rate of intergenic regions of both wild strains in Santeuil (μ_bs.HS_; unit: per site per year) and mutation accumulation lines in the laboratory (μ _bs.HS_; (Konrad *et al*. 2019); i.e., 1.82 × 10^-9^ per site per generation), we estimated the number of effective generations *C. elegans* reproduced in Santeuil per year (*N*_EG_; unit: generations per year) by:

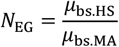

A Skygrid coalescent model was re-ran for HS2 using the 47 local non-recombinant strains and 6 external strains (Figure S4).

### Spatio-temporal patterns

Based on the GPS coordinates and notes recorded during field sampling, we mapped the distribution of different subclades (as defined on Figure 2) using ArcGIS v10.2 (ESRI, 2013). Euclidean distances between strains within each isotype were calculated (Tables S9-10). For each subclade, a Mantel test with 1,000 permutations was used to evaluate the correlation between Euclidean distance and sampling date difference among strains, keeping a single strain per sample (for example, JU1924 for S73). For subclades with significant Mantel test results, the slopes of regression lines can be interpreted as estimates of the dispersal rate of *C. elegans* (unit: meters per year) (Figure S5). For this analysis, the two remaining strains at the root of HS3.4 were conserved (JU1924 and JU4219). Pairwise Wilcoxon rank sum tests were used to assess whether significant spatial segregation exists among subclades, keeping a single strain per sample and not including the HS3.4 root strains (Table S8).

**Figure 2.**
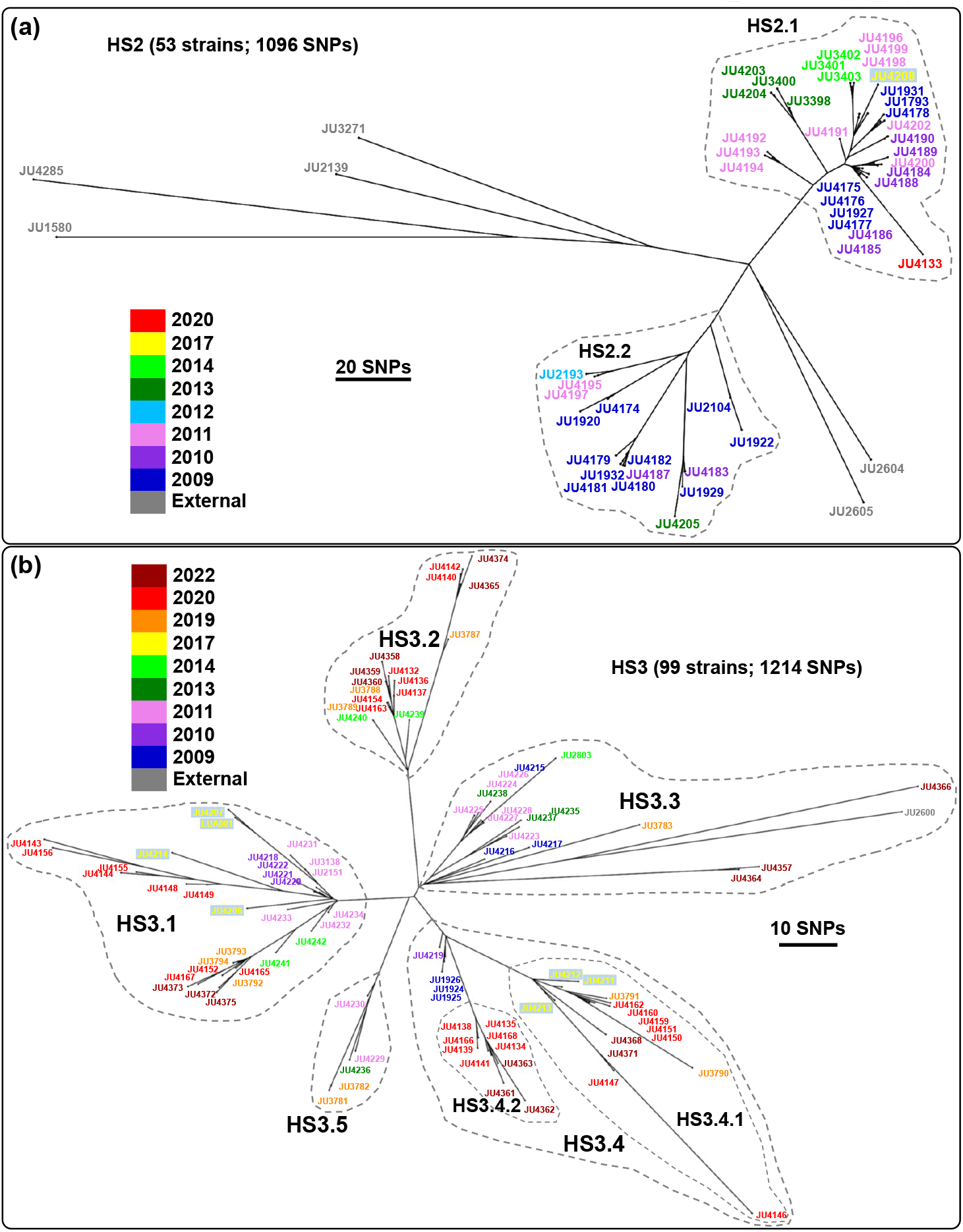
Haplotype networks of HS2 and HS3 strains. (a) HS2 genotypes. (b) HS3 genotypes. This figure includes strains from outside the Santeuil wood (i.e., external strains, in grey). In (a), to focus on mutation accumulation over time, the HS2 network is shown here without the three recombinants shown in Figure S3 of the Supplementary Materials. Subclades used for the spatial analysis are indicated by dashed-line polygons.

Each strain was assigned a one-dimensional coordinate on a simplified linear axis (Figure 6c), with the origin (0) set to the northernmost strain and values increasing southward (Table S1). This one-dimensional location was treated as a trait and used for visualizing the distribution and clustering of strains within each subclade on the MCC trees (Figure S6).

All statistical analyses were performed in R v4.1.2 (RCoreTeam 2023).

## Results

### Frequencies of co-occurring isotypes

The proportion of isotypes varied across years in our field collections (Scheirer-Hare rank test, year*isotype: *p*<0.003) (Figure 1). The HS1 isotype was predominant in 2009, then decreased and apparently disappeared in later years. The HS2 isotype became predominant, then HS3. Two new isotypes appeared in our collections in 2020 and 2022, which we named HS4 and HS5, respectively. These two isotypes are not mere recombinants of HS1-3, so immigration of new alleles likely occurred (Figure S2).

The HS2 isotype was found in several locations in France, at various distances up to 300 km away (Table S1). The HS3 isotype was only found outside the Santeuil wood a few hundred meters away in an apple (strain JU2600). The other three isotypes (HS1, HS4 and HS5) have so far not been found elsewhere.

### Local mutations and rare intra-isotype recombination

After the preliminary clustering in isotypes, we focused on intra-isotype polymorphisms, using whole-genome sequencing data of strains representing HS2 (n=56 strains) and HS3 (n=99 strains). As expected, the intra-isotype polymorphism level was markedly smaller than that among the local isotypes (Figure 1d). Although both indels and single-nucleotide polymorphisms (SNPs) contribute to genetic variation, in order to estimate mutation rates and calibrate them with other studies, we focused on SNPs, which are easier to call with high quality. For each isotype, we trimmed the genome of repeated and highly divergent regions to focus on genomic regions allowing high quality SNP calls. By distinguishing Santeuil wood strains (“local”) from other strains of the same isotype (“external”), we called a total of 1165 SNPs for 56 HS2 strains within the 63.07 Mb of the HS2 trimmed genome, in which 600 SNPs only occurred in six external strains, and 69 SNPs only occurred in three recombinants. After excluding both external and recombinant strains, 496 SNPs were found to be polymorphic among the 47 local HS2 strains. Similarly, a total of 1214 SNPs in 66.46 Mb were called for 99 HS3 strains. After excluding the single external strain (JU2600), 1147 SNPs were found in the 98 local HS3 strains.

From these polymorphisms, we analyzed the relationships between strains of a given isotype by building haplotype networks including both local and external strains (Figures 2 and S3). The trees in Figure 2 represent 1096 SNPs for HS2 and 1214 SNPs for HS3. For HS2, the external strains from outside Santeuil appeared at the same branching point. The positions of these external strains supported the idea that the mutations among Santeuil HS2 strains may have occurred locally in recent years; in other terms, these external strains can be considered as outgroups. In addition, most polymorphisms called among strains of each isotype were not found elsewhere in the CaeNDR strain set - the exceptions are likely due to convergent mutations or sequencing/mapping errors, especially from mononucleotide runs and segmental duplications that were not removed during trimming. Specifically, 87.9 and 85.5% of the polymorphisms within HS2 and HS3, respectively, were only found in the Santeuil wood. The detected polymorphisms did not cluster in a genomic region (Figure 3a), also consistent with them originating from new mutation rather than recombination. Finally, many of these polymorphisms could be found in several Santeuil strains, indicating that they appeared in the field and not subsequent laboratory culture. We conclude that the detected polymorphisms represent new, derived mutations occurring recently in the genomic background of this isotype. The mutational change could be polarized by inferring that the Santeuil private allele was derived.

**Figure 3.**
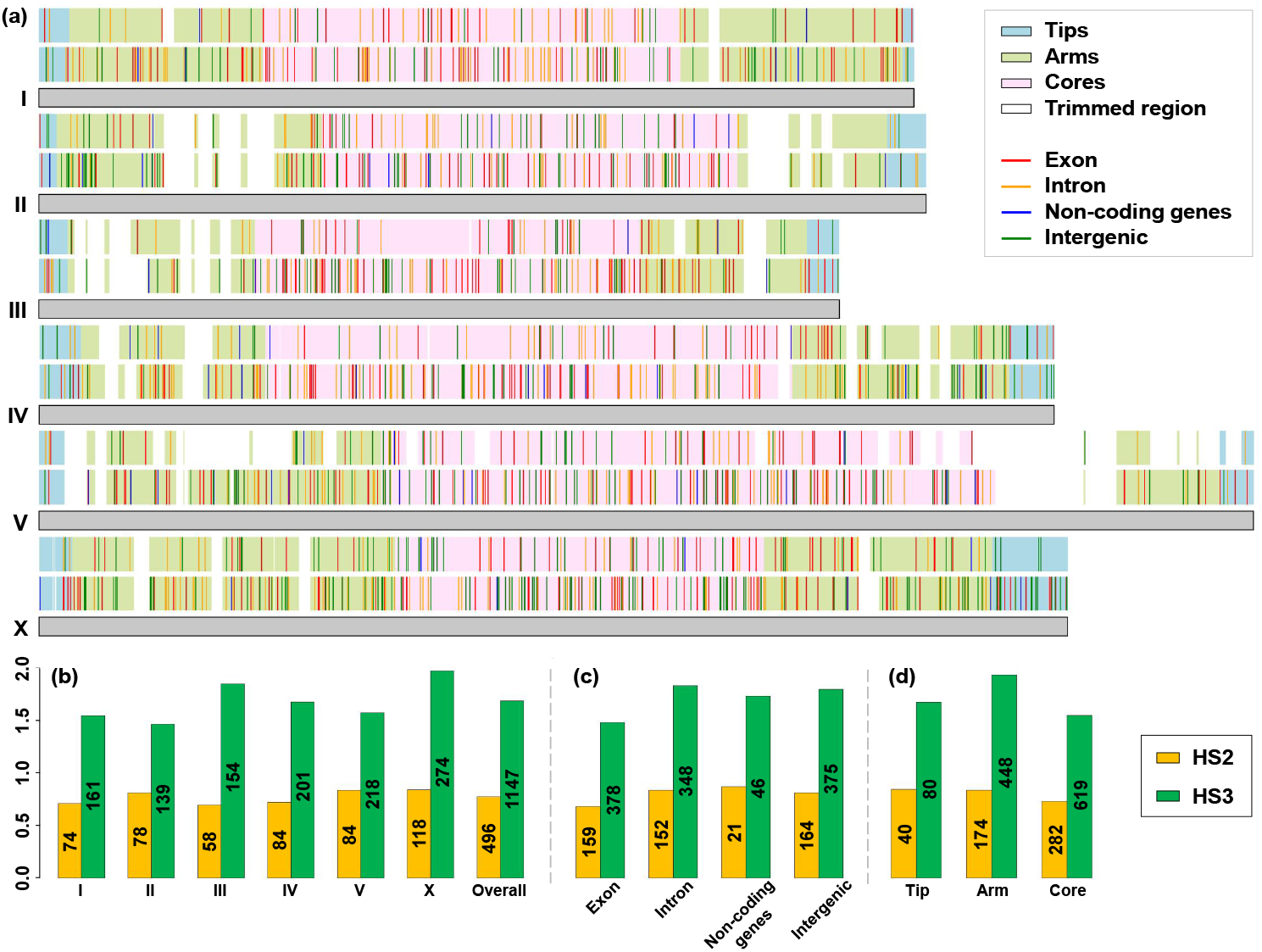
SNP distribution in different genomic regions. (a) SNP distribution along chromosomes in HS2 (upper) and HS3 (lower). (b-d) SNP density expressed in × 10^-5^ per site displayed by (b) chromosome, (c) genic region, and (d) recombination domain. The figure uses 496 and 1147 local SNPs for HS2 strains and HS3 strains, respectively. External strains and recombinants were not included. Blank regions are those removed in the trimmed genomes. In (b)-(d), the number of SNPs in each category is annotated on the corresponding bars. In (b), the distribution is not even among chromosomes (Chi-Square Goodness of Fit Test for even rates among chromosomes, *p* = 0.02 for HS3, *p* > 0.05 for HS2), with the X chromosome displaying the highest mutation rate. In (c), exons carry significantly fewer mutations per site than introns or intergenic regions (Chi-Square Goodness of Fit Test for genic regions, *p* = 0.02 for HS3, *p* > 0.05 for HS2). In (d), the boundaries of tips, arms, and cores refer to (Rockman and Kruglyak 2009). Arms carry significantly more mutations per site in HS3 (Chi-Square Goodness of Fit Test for tips, arms and cores, *p* = 0.002 for HS3, *p* > 0.05 for HS2). Detailed counts are in Table S5.

We then wondered whether outcrossing could be more frequent *within* isotypes than that previously estimated *between* isotypes, since outbreeding depression may be less relevant in the former case. For HS2, two recombination events occurred, as revealed by the diamond shapes in the network including all strains (Figure S3a,b), and concerned a total of three strains in our dataset. Specifically, with the SNP set that we could analyse, the JU3785 and JU3786 strains displayed whole-chromosome reassortments between the inferred parental genotypes, while JU3399 displayed one recombination event on four of the six chromosomes (Figure S3c,d). Each of these two events thus appeared consistent with a single outcrossing event. Removing the three recombinant strains resulted in a tree-like pattern, where the relationships between strains could be solely explained by mutation (Figure 2a). For the sake of simplicity, we excluded recombinants to infer mutational rate and pattern below. In the HS3 isotype, no recombination was detected or if so, only concerned so few mutations that this might correspond to the resolution of two recent mutations in the heterozygous state. We thus conclude that effective outcrossing events are also rare within these local isotypes.

### Mutational pattern

We analyzed the distribution of new mutations with respect to the six chromosomes, chromosome regions, gene annotation and nucleotide change. We performed these analyses on the local strains of the HS2 and HS3 genotypes separately (Figures 3 and 4). These analyses use 496 SNPs for HS2 and 1147 SNPs for HS3 and are thus more powered for the latter.

**Figure 4.**
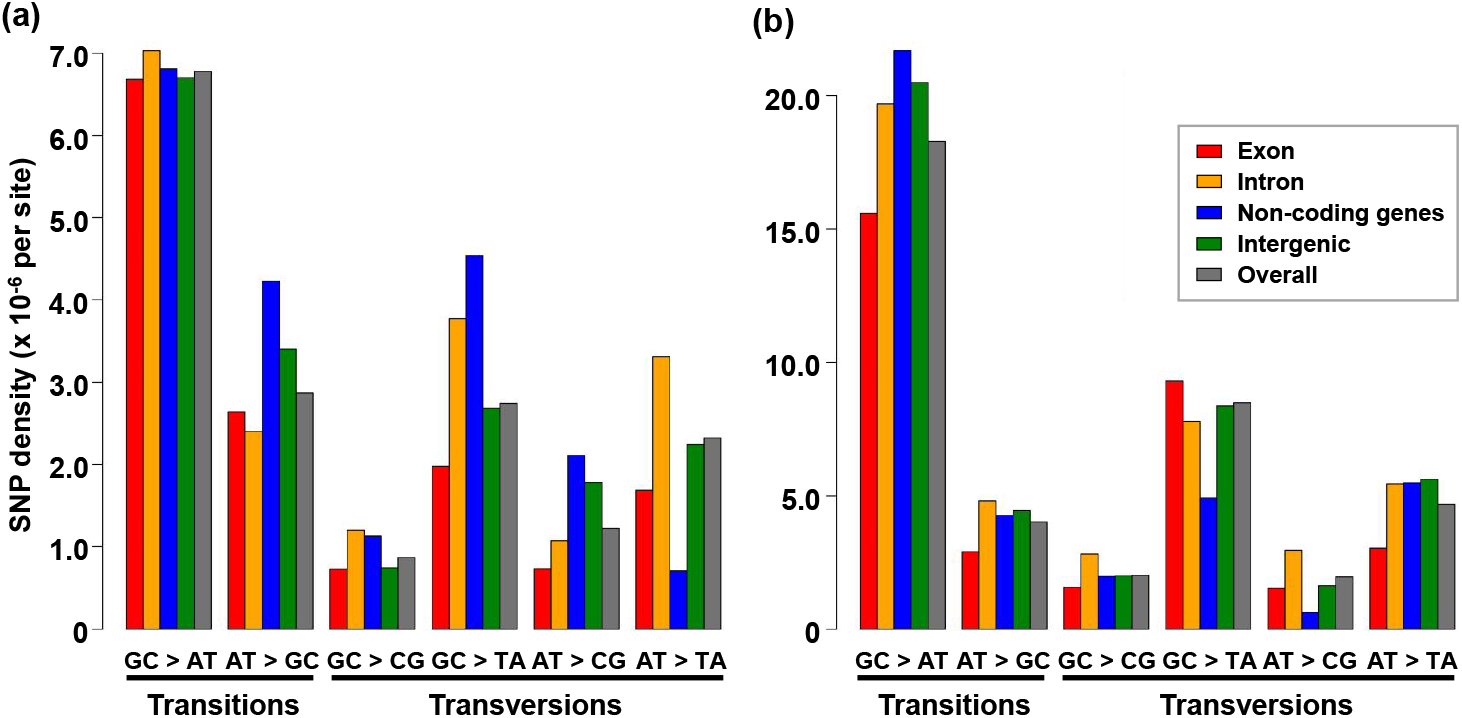
Mutational spectra expressed as density of different substitutions. (a) HS2. (b) HS3. The total number of mutations in each genic region category is indicated in Table 1 for each isotype. Detailed counts are in Table S6.

The distribution of mutations among the six chromosomes was uneven (Chi-Square Goodness of Fit Test for even rates among chromosomes: *p* = 0.02 for HS3, *p* > 0.05 for HS2). The X chromosome displayed the highest density, with 22% more substitutions per bp than autosomes for HS3, which is marginally significantly higher (Chi-Square test, *p*=0.055) (Table S5.

**Table 1.**
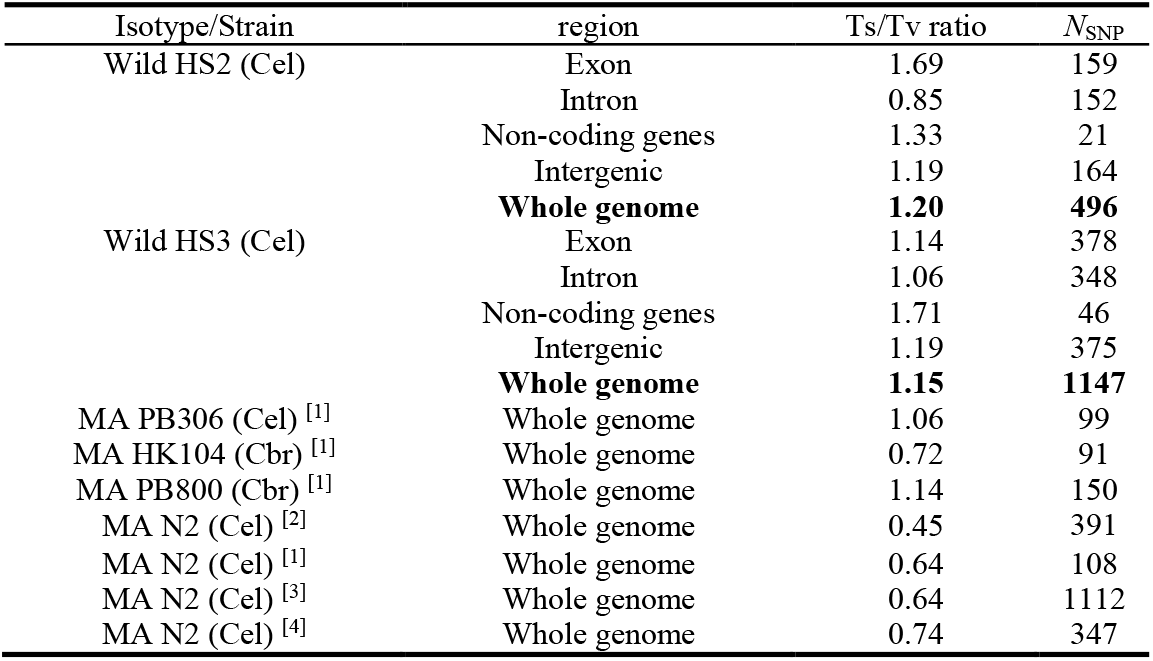
Ts/Tv ratios of different genic regions in wild isotypes sampled in the field (i.e., HS2 and HS3), and Ts/Tv ratios of the whole genome from mutation accumulation (MA) lines cultured in laboratory from ^[1]^(Denver *et al*. 2012), ^[2]^(Denver *et al*. 2009), ^[3]^(Konrad *et al*. 2019), and ^[4]^(Saxena *et al*. 2019). Cel - *Caenorhabditis elegans*; Cbr - *C. briggsae. N*_SNP_ is the number of SNPs in each category.

We then partitioned each chromosome in core, arms, and tips, following recombination regions from (Rockman and Kruglyak 2009), where arms are defined by a higher recombination rate. The core (central) region of *C. elegans* holocentric chromosomes recombines at a lower rate and harbors fewer polymorphisms on average at the species scale, fewer repeated elements, and more conserved and essential genes (Consortium 1998; Fraser *et al*. 2000; Rockman and Kruglyak 2009; Andersen *et al*. 2012; Crombie *et al*. 2019). In the HS3 dataset, arms carried significantly more mutations per site than other chromosomal regions (Figure 3d; Chi-Square Goodness of Fit Test for tips, arms and core: *p* = 0.002 for HS3, *p* > 0.05 for HS2, although with the same trend).

We then partitioned the SNPs in different genic regions (exons, introns, intergenic, non-coding gene). Exons displayed a significantly lower mutation density and substitution rate than the other genic regions (Chi-Square Goodness of Fit Test for density in different genic regions: *p* = 0.02 for HS3, with the same trend in HS2) (Figure 3c, Table S5). This result is consistent with purifying selection on exons.

To investigate the higher SNP density on chromosome X in HS3, we further partitioned autosomes and X chromosome by genic regions (Table S5). For HS3, the SNP density of intergenic regions was not significantly different on the X chromosome versus autosomes (Chi-Square test, *p* = 0.3); idem for introns (*p*= 0.9). In contrast, SNP density on exons was higher on the X chromosome than on autosomes (*p* = 0.025), suggesting weaker purifying selection.

We could polarize each mutation and thus classify single-nucleotide substitutions in six chemical categories (Figure 4). GC to AT transitions were particularly frequent in both isotypes, suggestive of spontaneous deamination of cytosines (Lindahl 1993) (Figure 4). The expected ratio of transitions to transversions (Ts/Tv) should be 0.5 if the mutation pattern were random. In our dataset, transitions were found more frequently than transversions, with a Ts/Tv of 1.20 for HS2 and 1.15 for HS3 (Table 1). At this whole-genome level, the two isotypes did not exhibit significant differences (Chi-Square test, *p* > 0.05). In HS3, the Ts/Tv ratio was noticeably lower in intronic regions (Table 1).

The *C. elegans* genome contains less G/C than A/T bases (36% CG), although this trend is less strong in exons (Table S5). We weighted the six substitution rates with the GC content of each genic region category. The mutational pattern resulted in a significant bias towards AT in exons of HS3 (Chi-Square test, *p* < 0.001), but not significant in HS2 (Table S6). To assess the effect of mutation on GC content evolution, we estimated the relative proportion of mutations from GC to AT or TA over those from AT to GC or CG. Weighting the number of relevant substitutions in each genic region, we found ratios of 1.75 and 0.85 for HS3 and HS2, respectively, thus a net mutational bias towards a higher AT content in HS3. However, we note the difference between the two isotypes.

Finally, using coding gene annotations, we estimated SNP density of non-synonymous versus synonymous mutations within coding sequences (Table S7). We found ratios of 2.3 and 3.0 for HS2 and HS3, respectively. We did not detect a significantly different ratio in the Santeuil sets compared to the mutation accumulation lines (PB306 from (Denver *et al*. 2009) and N2 from (Denver *et al*. 2009; Denver *et al*. 2012), and (Konrad *et al*. 2019); Chi-Square test, *p* > 0.05).

### Temporal signal: mutation rate and number of generations per year

The branching pattern over time differed for strains of the two isotypes. For HS2, the Santeuil wood lineages diverged from the external strains along two branches that could correspond to two independent immigration events or to local mutation and selection trimming away branches (Figure 2a). By contrast, the HS3 isotype strains showed a star-like relationship, consistent with a single founder and lineage expansion along the years (Figure 2b). A distinct temporal signal can be seen: the strains collected in earlier years are on inner nodes of the phylogeny, while recently collected ones are on the outside (Figure 2). The temporal pattern was particularly clear for HS3 and appeared weaker for HS2, in part because of its decline.

Using sample collection dates, we inferred the most credible evolutionary scenario using Bayesian Maximum Clade Credibility (MCC) trees, for HS2 and HS3 separately (Figure 5). Given the lack of detected recombination with this dataset, the topology of the tree did not carry ambiguities, while the dates of nodes were the best estimates. The two main subclades of the HS2 tree branched around year 1990 while the HS3 tree was rooted around year 2000. When adding the external strains of HS2, a possible isotype root was estimated to have occurred before 1960 (Figure S4).

**Figure 5.**
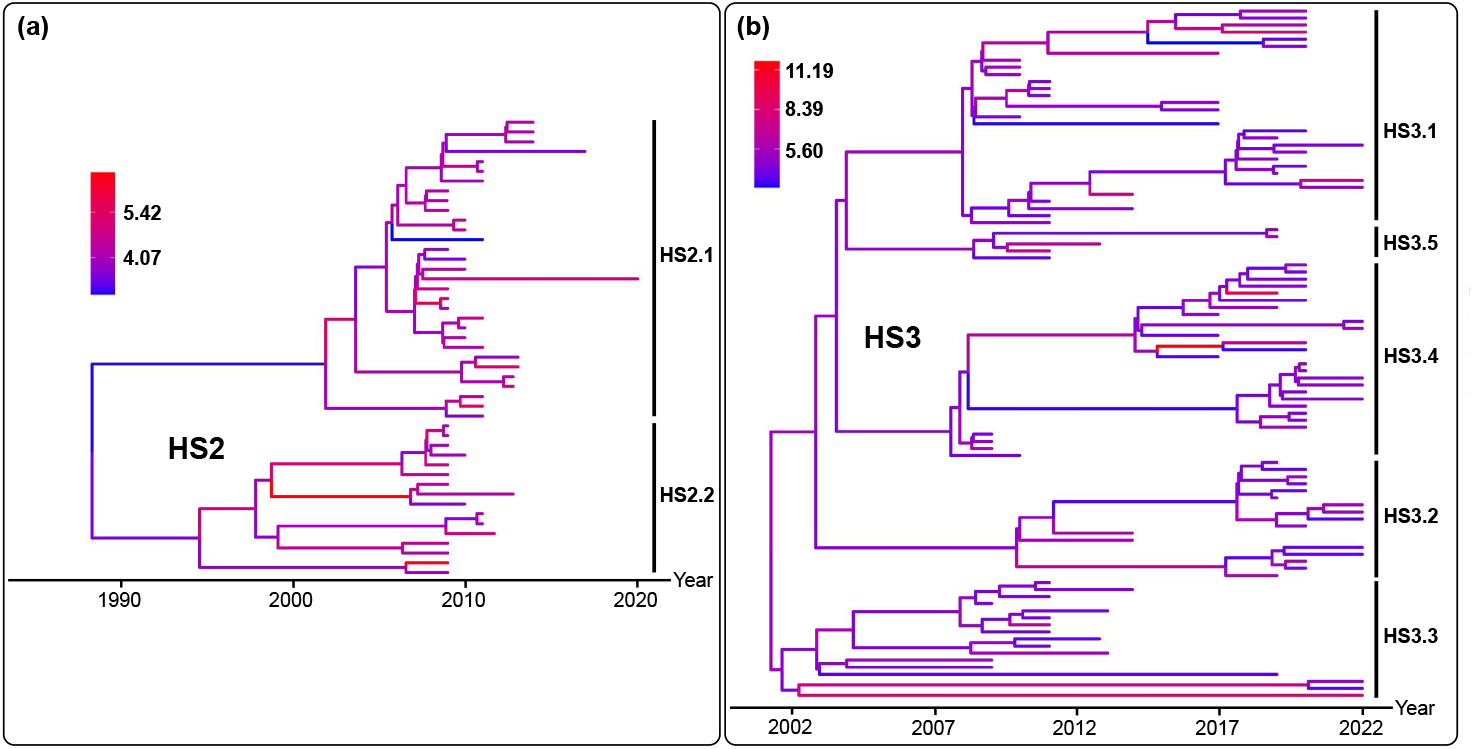
Maximum Clade Credibility trees of Bayesian Skygrid demographic model with tip-calibration. (a) HS2 genotypes. (b) HS3 genotypes. Branch color represents substitution rates in intergenic regions (× 10^-8^ per site per year). Note the different scales in the two panels.

**Figure 6.**
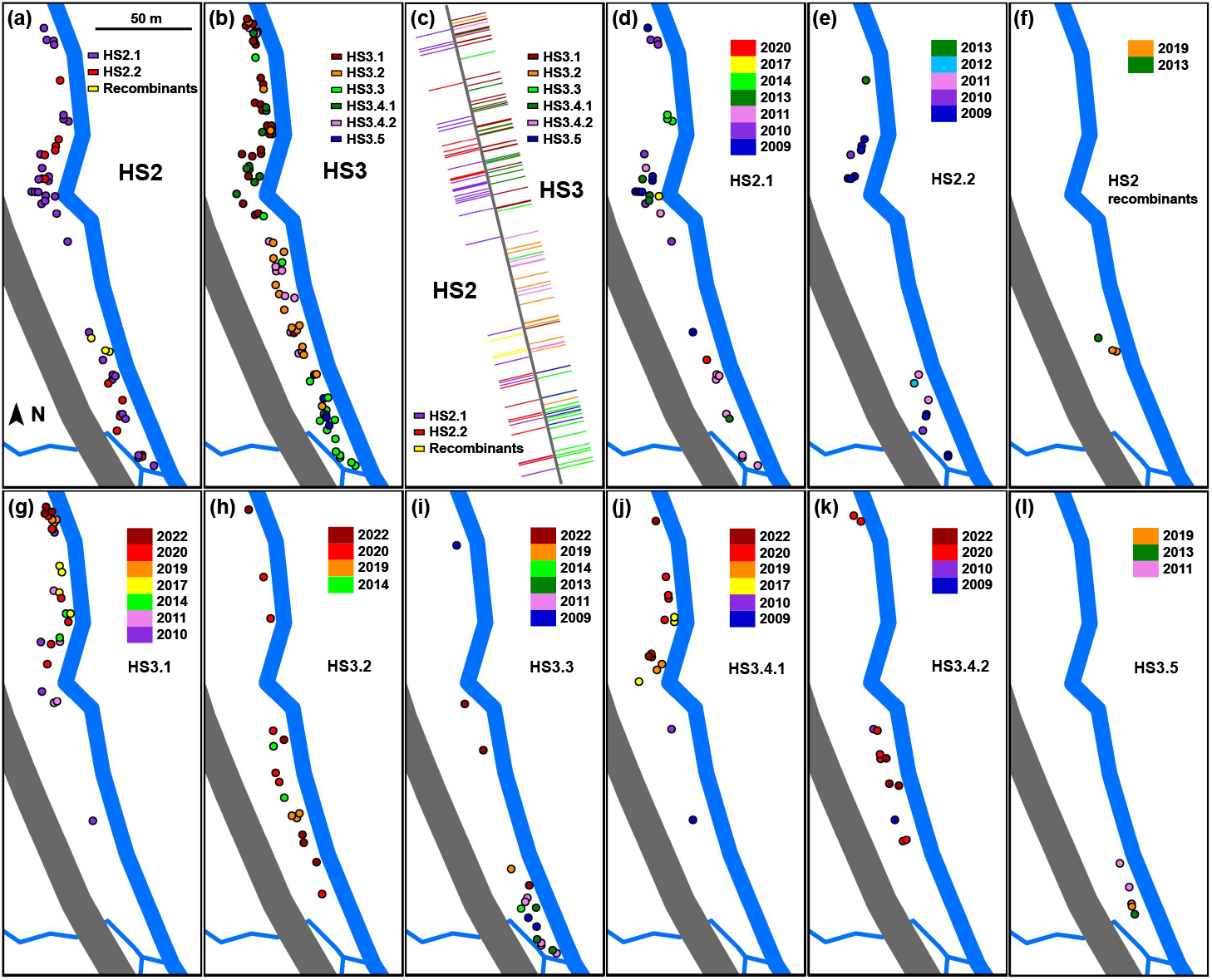
Spatial distribution of the isotypes and subclades across years. (a) Spatial distribution of subclades within HS2. (b) Spatial distribution of subclades within HS3. (c) Spatial distributions projected along a simplified straight line. In (a)-(c), the colors indicate the subclades. (d-l) Spatiotemporal distribution of each subclade. Detailed information about strain clustering in subclades is indicated in Figure 2. In (d)-(l), the colors indicate sampling year of each strain. The four strains in the basal position of clade HS3.4 are shown in plots of both subclades HS3.4.1 (j) and HS3.4.2 (k). The gray line represents a railway and the blue lines represent water bodies, based on data from OpenStreetMap (2021).

Based on the MCC trees built using local non-recombinant strains, we estimated the average number of substitutions per year for each isotype. In order to exclude mutations that could have occurred in the laboratory, we removed the branches at the tree tips. In addition, we selected intergenic regions, as the rate per site per year was significantly lower in exons (see above, Figure 3c), which are more likely to be under purifying selection. In intergenic regions, mutation rates for the HS2 and HS3 natural populations were estimated to 4.13 and 4.92×10^-8^ mutations per site per year, respectively, with overlapping confidence intervals (Table 2). With a 100 Mb genome, this corresponds to 4 to 5 mutations per haploid genome per year.

**Table 2.** Substitution rates (*μ*_bs_; × 10^-8^ per site per year) in exon, intron, non-coding genes, and intergenic regions, and number of effective generations per year (*N*_EG_). *N*_SNP_ is the number of SNPs in each category. † The *μ*_bs_ of intergenic regions was employed to estimate *N*_EG_ referring to *μ*_bs_ of intergenic regions of mutation accumulation lines (i.e., per site per generation) from (Konrad *et al*. 2019). Intervals in brackets represent 95% highest posterior density.

|  | HS2 |  | HS3 |  |
| --- | --- | --- | --- | --- |
| | $N_{\text{SNP}}$ | $\mu_{\text{bs}}$ | $N_{\text{SNP}}$ | $\mu_{\text{bs}}$ |
| Exon | 159 | 3.44 (2.34 - 4.54) | 378 | 3.64 (3.04 - 4.26) |
| Intron | 152 | 4.30 (2.98 - 5.75) | 348 | 4.43 (3.67 - 5.20) |
| Non-coding genes | 21 | 4.13 (1.74 - 6.85) | 46 | 4.80 (3.20 - 6.40) |
| Intergenic<sup>†</sup> | 164 | 4.13 (2.84 - 5.53) | 375 | 4.92 (4.14 - 5.69) |
| $N_{\text{EG}}$ | 22.69 (15.60 - 30.39) | | 27.03 (22.58 - 31.26) | |

To convert this substitution rate in number of generations per year, we used the spontaneous mutation rate per generation measured using laboratory mutation accumulation lines. Two recent independent studies gave similar results (Konrad *et al*. 2019; Saxena *et al*. 2019). Based on the rate of 1.82 × 10^-9^ per site per generation in intergenic regions from (Konrad *et al*. 2019), we estimated the number of effective generations in Santeuil natural populations to 22.7 (15.6 - 30.4) and 27.0 (22.6 - 31.3) generations per year for HS2 and HS3, respectively.

### Spatial structure within the Santeuil field site

We then used the new mutations detected above to study spatial structure and dynamics in the field. Because the trees in Figure 2 represent actual animal lineages across generations, the fate of mutations at the root of each clade could be followed in space - bearing in mind the spare sampling of our collection compared to the actual population size.

We first plotted the data on two-dimensional spatial maps (Figure 6), subdividing each isotype into main subclades: the HS2 isotype into two main subclades plus three recombinants, and the HS3 isotype into six main subclades, as shown in Figures 2 and S5. For HS2, the two subclades were found from North to South and were not significantly spatially segregated from each other (Wilcoxon Rank Sum Test, *p*>0.05). Strikingly, the three strains deriving from the two independent recombination events were found in a central region of the transect almost devoid of other HS2 strains - however, this corresponded to few cases. In contrast, for the six main HS3 subclades, which branched more recently than the two HS2 main subclades, we found highly significant segregation, for example of HS3.1 (North) with HS3.3 or HS3.5 (South) (Figure 6, Table S8).

We further asked whether we could detect a spatio-temporal signal within each subclade. We reasoned that if a new mutation occurred at one end (or even outside) of the transect, we may detect a spatio-temporal signal of dispersal. Mantel tests between pairwise Euclidean distance of strains and their sampling date distance found that two of eight tested HS3 subclades showed a significant signal of directional migration (Figure S5), both from South to North. The regression slopes can be interpreted as estimates of the dispersal rate of the *C. elegans* lineage carrying the mutations at the root. The subclade showing the strongest signal, HS3.4.1, gave a slope of 6 meters per year (C.I. 4.8-7.2), which if sampling was sufficient, is a minimal estimate of the maximal dispersal speed.

To represent the spatial data including sampling year and all genetic relationships, we simplified the geographical coordinates as a one-dimensional projection on the transect line (Figure 6c) and colored the tips of the phylogenetic tree by the spatial position of the strain (Figure S6). Clustering of spatial positions on the phylogenetic tree is apparent, including across years. The broad patterns of the HS3 subclades are visible: for example, HS3.1 strains in a Northern position (smaller coordinate values) and HS3.3 and HS3.5 in a Southern position. In addition, smaller subclades within HS2 and HS3 show clustering, with the caveat of limited sampling.

Clustering of different subclades was only apparent because movement across the transect occurred. Focusing on the early years of the HS3 tree, the same mutation at the root of the sister subclades HS3.1 and HS3.5 was found in strains collected on opposite transect sides, implying movement across 200 meters at the root or early history of one or both subclades. Compared to the HS3.1 strains located in the North, early-year strains of basally branching subclades were almost all found in the Southern half of the transect, suggesting a northwards movement in the HS3.1 root, even prior to within-clade movement of Figure S6. Alternatively, this among-subclade pattern may be explained by repeated southwards movement. At a fine temporal scale, some closely related pairs of strains, for example JU4135/JU4168, JU4138/JU4166 or JU3788/JU4154 show evidence of large movement (Figure S6). These 1 mm long nematodes can move by themselves, yet faster dispersal vectors are larger invertebrate carriers (BarriÈre and FÉlix 2005; Caswell-Chen *et al*. 2005; BarriÈre and FÉlix 2007; FÉlix and Duveau 2012; Petersen *et al*. 2015). We wondered whether these animals carried C. *elegans* with the same genotype as the stem on which they were collected, that is whether, in this stem, *C. elegans* may have disembarked from or embarked onto this larger invertebrate (Tables S1, S11). We found one case of concordance between the *C. elegans* genotype on the animal and the plant, six cases of discordance, and one mixed case where two different carriers gave contrasted results (Table S11). A limitation of this analysis is that a plant substrate itself - or an animal - often carries different *C. elegans* genotypes (Richaud *et al*. 2018).

In summary, despite the fact that migration occurred, the mutated alleles at the root of these subclades did not all spread across the 300 m-long field site within 15-20 years, giving a bound on dispersal at this scale. On the other side of the coin, our observations imply cases of movement across the transect in the span of one or few years.

## Discussion

### Number of nucleotide substitutions and generations per year

In this study, we estimated in a *C. elegans* natural population a nucleotide substitution rate of 4 to 5×10^-8^ mutations per base pair per year. Previous studies that time-calibrated the species tree estimated the age of tree nodes in number of generations (Cutter 2008; Fusca *et al*. 2025; Picao-Osorio *et al*. 2025). However, our estimate of substitution rate per year now suffices to calibrate species distance on the genus phylogeny, independently of the mutation rate estimates in laboratory mutation accumulation lines and the estimated number of generations per year.

Although we do not need to scale the natural mutation rate to the laboratory mutation rate, the former will depend on the number of effective generations per year. For example, the estimates of mutation rate are somewhat lower for HS2 than for HS3 (yet with overlapping credible intervals); a lineage remaining at low frequency, such as HS2, could undergo fewer effective generations and mutations per year than a then thriving lineage such as HS3. The extrapolation of this local measure should thus be taken with caution, as demography, temperature and duration of the lifecycle will also vary with the species. For example, tropical species may see a higher number of generations per years; conversely, some species develop slower than *C. elegans*, at least in laboratory conditions, for example *C. inopinata* (Kanzaki *et al*. 2018) or more distant species in the genus (Stevens *et al*. 2019). Interestingly, Fusca et al. (2025) found low variation in the substitution rate among evolutionary branches of the *Caenorhabditis* genus, which makes our estimate potentially robust. Therefore, although further measures in different populations and species would be welcome, using this temperate *C. elegans* population appears a useful first estimate of the substitution rate per year.

When calibrated on the laboratory mutation rate, the substitution rate translates into 15 to 30 effective generations per year, a key parameter of *C. elegans* natural ecology. Thus, surviving lineages in these populations displayed an average generation time of about two weeks, which is several-fold higher than the minimal generation time of three days in favorable laboratory conditions.

Two factors likely contribute to a longer lifecycle in natural populations. First, the duration of direct developmental cycles is strongly affected by environmental factors, such as temperature, food and pathogens. For example, development can last over a week at temperatures below 15°C, often encountered in the fall in Santeuil. Second, resources are limited, resulting in developmental arrest of individuals, including dauer diapause. In individual lineages, generations with fast direct development alternate with generations developing via a dauer stage. Given the observed census size and developmental stage composition of populations, dauer induction at high population density and low food, and the patchiness of suitable substrates, it is probable that no more than two to three generations of direct development occur before an individual lineage is induced to dauer (FÉlix and Duveau 2012). Diapause can last for months when resources are scarce.

In temperate areas like Santeuil, the generation time likely varies seasonally, with two consequences on *C. elegans* generation time: 1) the availability of food patches peaks in the fall with the bulk of plant decomposition. 2) development slows down progressively during the fall when temperatures decrease. We previously found that, in an apple orchard in Orsay, near Paris, France, *C. elegans* proliferates in the fall and not summer and winter (FÉlix and Duveau 2012); a fall peak of *C. elegans* was also detected in compost heaps in Germany by (Petersen *et al*. 2014). In Santeuil, we collected *C. elegans* in the fall when decomposing stems are easier to find. We note that the Santeuil wood undergrowth is more shaded and humid than the Orsay orchard apples, thus cooler in summer, and may harbor *C. elegans* populations for a longer time. We indeed found the species in summer at the two timepoints when we collected (06 Jul 2012, 07 Aug 2013) (Richaud *et al*. 2018) (Table S1). In summary, *C. elegans* development may occur fast when conditions are optimal in late summer/early fall, then temperatures decrease in the fall and the generation time likely slows down. Given the diapause arrests and the environmental modulation of the direct lifecycle, our estimate of 15 - 30 generations per year appears consistent with the range of expectations. Some lineages may undergo fewer generations than others in a given year, but the likelihood that they contribute to the next generations would be smaller.

A limitation concerning the conversion of mutation to generation numbers is that the mutation rate in the laboratory may not match that in the wild. We used estimates from the typical laboratory environment at 20°C, with direct development, and the rate may vary with the environment. Nevertheless, by testing the effect of an oxidative environment, *a priori* expected to contribute to mutation, (Rajaei *et al*. 2021) found that mutation rate and pattern were not strongly affected. In addition, the mutation rate could differ among genetic backgrounds. However, Denver et al. (2012) did not report a large variation among *Caenorhabditis* strains. Finally, while the strength of purifying selection may differ between natural populations and laboratory mutation accumulation lines, this difference has limited impact on our estimation given that we only used intergenic regions where selection is expected to be weaker than on exons, thus minimizing potential bias.

### Genomic mutation pattern and detection of selection

We compared the mutation and SNP density pattern to i) a null expectation with no mutational bias; ii) that observed in laboratory mutation lines, where selection is minimal; iii) in some cases the species-wide pattern.

We found a higher SNP density on chromosome arms compared to centers and tips, corresponding to regions where recombination is highest. This mutation pattern is similar to that detected in laboratory mutation accumulation lines (Konrad *et al*. 2019; Saxena *et al*. 2019) and thus consistent with meiotic events of recombination being mutagenic. Alternatively, as suggested by Konrad et al. (2019), this correlation may be primarily explained by the differential abundance of mutagenic motifs. An alternative selective explanation is possible, essential genes being enriched in central regions of *C. elegans* chromosomes (Kamath *et al*. 2003).

More surprisingly, we detected a significantly higher SNP density on the X chromosome compared to the autosomes, albeit of modest size (Table S5). This pattern contrasts with 1) the pattern in laboratory mutation accumulation lines from the N2 strain (Konrad *et al*. 2019), where the sex chromosome had the lowest rate of substitutions; 2) the species-wide pattern where chromosome X carries only 14.1% of the polymorphisms (CaeNDR version 20250625, using hard-filtered VCF), whereas 23.9% of our intra-isotype polymorphisms are on chromosome X. The different chromatin context of the X chromosome compared to autosomes (Kelly *et al*. 2002) may modulate the mutation rate but cannot explain the results in these three conditions. Moreover, the GC nucleotide content of the X chromosome is not higher than that of the autosomes (Table S5). The fast rate of X evolution in Santeuil may be due to the genetic background: the *C. elegans* PB306 strain showed a marginally significantly higher mutation rate on the X chromosome compared to autosomes, whereas the N2 reference strain did not (Denver *et al*. 2012). Alternatively, the fast rate of X evolution in the natural populations could be due to a population genetic effect (selection), such as more frequent purifying selection on autosomal exons. In line with this hypothesis, the proportion of nucleotides in exons is of 34.4 to 41.9% on autosomes but of 29.0% on the X chromosome, and essential genes are depleted on the latter (Kamath *et al*. 2003). In addition, the density of SNPs within exons was higher on the X chromosome than on autosomes, suggesting that purifying selection is stronger on autosomal exons. An alternative explanation for the “faster X” evolution at larger timescale is positive selection of new mutations - however, this seems unlikely to affect patterns at the scale studied here. In conclusion, the slightly higher SNP density on the X chromosome appears to derive from a lower proportion of exons allied with a higher SNP density on exons on the X chromosome compared to autosomes.

A possible sign of purifying selection in our data is indeed the depletion of substitutions in exons, over the whole genome. The SNP density in exons is 20 and 23% lower than in intergenic regions and introns, respectively, which cannot be explained by the GC content of exons. Consistent with our findings, Exposito-Alonso et al. (2018) also found that purifying selection acts pervasively on coding regions in natural populations of *A. thaliana*. This demonstrates the ubiquity of purifying selection in natural populations of selfers, although its strength may depend on environment and population history. This pattern resembles that in laboratory mutation accumulation lines, where selection is minimal yet present (Konrad *et al*. 2019; Saxena *et al*. 2019). That this results from selection is tantalizing, but alternative hypotheses related to nucleotide context, chromatin structure, transcription and DNA repair mechanisms are possible. Within exons, we did not detect a difference in the ratio of non-synonymous to synonymous mutations compared to laboratory mutation accumulation lines, likely due to a lack of statistical power at this scale.

The transition-to-transversion ratio in the local natural populations is almost double to that in mutation accumulation lines in the N2 reference background, more similar to those in the PB306 background (Table 1), and close to species-wide polymorphisms (1.29; CaeNDR 20250625, using hard-filtered VCF). The higher rate of transitions may derive from the mutational pattern that favors chemical conservation of purines and pyrimidines, for example through the deamination of cytosine to thymine. Alternatively, the difference between our dataset and mutation accumulation lines could be explained by selection favoring nucleosome conservation or, in exons, synonymous or conservative amino-acid changes that are favored by transitions (Babbitt and Cotter 2011). Finally, as the statistical difference between HS2 and HS3 mutational patterns exemplifies, the genetic background and environment contexts may influence this mutational pattern as well.

In summary, the estimated 500-generation evolutionary timescale on a selfing metapopulation may allow the detection of purifying selection on new mutations at the genome scale, yet the signal is weak if any. Particular contrasts with the laboratory mutation accumulation lines are the higher mutation rate on the X chromosome (both isotypes) and the high Ts/Tv ratio for the HS3 isotype. Studies of mutation using the Santeuil genetic backgrounds in the laboratory may help in the future to determine whether this is due to the different genetic background; alternatively, the natural environment, or selection, may influence these mutational patterns.

### Spatial structure and isotype turnover

This study along a 300 m field site could detect a spatial structure of new mutations across 14 years, thus providing a picture of eco-evolutionary processes at a fine spatio-temporal scale.

*C. elegans* proliferates in patchy bacteria-rich decomposing plant matter. Large populations (over 100,000 individuals) always include development of young larvae into the dauer diapause stage, which forms the basis for a boom-and-bust dynamic (FÉlix and Duveau 2012; Richaud *et al*. 2018). In the Santeuil field site, herbaceous plant stems (10 cm to 1 meter) start decomposition at their base and fall in succession, chiefly in autumn, covering the ground in a discontinuous manner. Therefore, the density distribution of *C. elegans* is highly heterogeneous across the site at a given timepoint. Compared to the well-studied and larger meadow patches of the Glanville fritillary butterfly in Finland (Hanski 2011; Multigner *et al*. 2025), these *C. elegans* patches are smaller and ephemeral. Across the field site, differences are visible, with a higher density of decomposing plants at the North and South ends compared to the middle and a different plant composition: for example, the coltsfoot *Tussilago farfara* was only found on the North side across all sampled years. Finally, at a larger scale, the Viosne stream bank is a stable habitat patch over these years, more similar to a Glanville fritillary butterfly meadow patch. Thus, metapopulation structure may occur at several spatio-temporal scales for *C. elegans*.

In this context, dispersal is a key parameter for population dynamics. Our mutation tracking provides a spatio-temporal scale at which *C. elegans* do not freely disperse. Specifically, the mutations at the root of the different HS3 subclades, estimated to have occurred between 2002 and 2008, have not all spread through the whole 300 m transect. This lack of spatial homogenization is compatible with a dispersal rate of few meters per year as estimated for HS3.4.1. Field sampling of a microscopic animal is inherently limited given the large population census size, which alters our ability for accurate dispersal estimates. Overall, our results thus show that *C. elegans* dispersal is limited at scales of 100 m over ten years.

The structure observed when considering new mutations within an isotype is likely mostly mediated by limited dispersal capacity, rather than differential selection. *C. elegans* dispersal can occur via both active self-movement and transport by animal vectors. Self-movement can be quantified in the laboratory on a surface (Husson *et al*. 2013) or in a volume (Kwon *et al*. 2015). *C. elegans* adults can move at an instantaneous speed of 0.2 mm/s on surfaces without food (Ramot *et al*. 2008). Extrapolating this instantaneous speed would allow movement over 20 m per day - as unlikely as Howard Berg’s imaginary molecule moving through a classroom in one second according to its instantaneous Brown motion speed (Berg 1993). *C. elegans* indeed uses biased random walks, even when chemotacting towards an odour, with turns and bouts of backward movement (Pierce-Shimomura *et al*. 1999; Roberts *et al*. 2016) analogous to runs and tumbles of *E. coli* (Berg and Brown 1972). *C. elegans* diffusion coefficient on smooth agar surfaces is on the order of 0.01 mm^2^/s (Helms *et al*. 2019). In real life, the animals slow down on food and, across the reproduction cycle, larvae are slower than adults; further delays occur in the three-dimensional natural environment, with liquid volumes where swimming is inefficient, high surface complexity, surface tension, stickiness, bacterial biofilms affecting movement (Darby *et al*. 2007). How far a single animal - during its lifetime until reproduction - would move remains unclear, but likely does not exceed a few decimeters for long-lived dauer larvae and centimeters for direct developers. At these short distances, nematodes can move in a random or semi-directional manner within a substrate and possibly between adjacent plant stems.

Longer-range dispersal of *C. elegans* may be mediated by physical causes such as rain, wind, and here the waterstream along the field site. Concerning the latter, fast movement appears to often run northwards, in opposite direction to the stream, such as mentioned for HS3.1 (root and within-clade dispersal) and also exemplified by single strains with a Northern coordinate within a Southern clade (Figure S6). Alternatively, slugs, snails, isopods and insects are able to disperse *C. elegans* individuals at medium-range distances (meters) (BarriÈre and FÉlix 2005; Caswell-Chen *et al*. 2005; FÉlix and Duveau 2012; Petersen *et al*. 2015; Schulenburg and FÉlix 2017; Fialova and Tuf 2025). Maximal movement speed was found in the JU4135-JU4168 pair of the HS3.4.2 lineage (Figure S6): the two strains were found at a distance of 173 meters and did not display any SNP divergence in our dataset, thus, with an average substitution rate of 5 mutations per year, diverged for less than a year. We caution that this maximal speed is sensitive to possible strain errors; however, the occurrence of several similar cases indicates occasional faster movement in an overall static population. Likely less frequent contributors to longer-range movement are vertebrates such as birds and mammals (including humans), which could move *C. elegans* within the site or further (kilometers). For example, the HS2/JU1793 isotype was found at various distances up to 300 km with a divergence date estimated to 50 years (Figure S4). Strains of the same isotype can be found on different continents, albeit rarely (Andersen *et al*. 2012; Lee *et al*. 2021): the “MY23” isotype appears to be a dispersal record holder, having dispersed in recent years across three continents (South America, Europe, and East Asia). The present threshold for *C. elegans* strains to be called of the same isotype at CaeNDR corresponds to about 1000 SNPs, thus 100 years of mutation on each branch (if the strains were sampled around the same years and no recombination occurred). As in the present study, tracking of recent mutations is a convenient approach for assessing dispersal in natural populations of microscopic organisms and will be of great value for future eco-evolutionary studies.

While the spatial structure within an isotype is largely mediated by dispersal ability, different isotypes and their spatio-temporal distribution could be affected by selection. During the 14-year period, we observed a turnover of isotypes with immigration of new alleles. One isotype (HS1) faded away early in our samples. The two HS2 local subclades branched around year 1990, which also corresponded to their divergence with external strains, and faded in the late years of our study. The local HS3 tree was rooted around year 2000 (Figure 5); given the lack of a clear outgroup, the immigration of the first individuals in Santeuil could have preceded this date. HS4 appeared in our samples in 2020, and HS5 in 2022. We cannot rule out that these isotypes were proliferating locally at low frequency in the early years, yet we find it unlikely that the larvae could subsist for several years as a “dauer bank” (by analogy with seed banks of plants). Therefore, the Santeuil wood appears to harbor a succession of selfing isotypes, some of them with a residence time of at least 20 years, and co-occurrence of a few isotypes at a given time. Neutral processes could contribute to isotype distribution and turnover dynamics, for example through limited dispersal and population bottlenecks favoring stochastic losses. Alternatively, micro-habitat heterogeneity in parameters such as microbial community, predators or physicochemical properties could result in differential selection. Laboratory competition experiments have demonstrated that diversifying selection, for example on resistance to pathogens or other stresses, may help maintain several local isotypes on timescales of a few generations (Richaud *et al*. 2018). At a larger scale, how intra- and inter-annual environmental variables, such as fluctuations in temperature, precipitation, or pathogen prevalence, affect the persistence of local isotypes remains to be investigated.

In conclusion, the present study provides an exemplary case for eco-evolutionary research of nematode natural populations at a fine scale. Studying metapopulations of these small animals requires the consideration of different spatio-temporal scales, dominated by different processes. Studies at a larger spatial scale may help understand patterns of recombination that lead to isotype diversity and thus complement the present study of local mutational patterns.

## Data availability statement

Raw sequence reads are available at CaeNDR.org or deposited at NCBI under project number PRJNA1280162.

## Acknowledgements

We are grateful to Gaotian Zhang and Joëlle Barido-Sottani for help and discussions on data analysis. We thank Asher Cutter and Amanda Gibson for constructive comments on the manuscript. We thank Buck Samuel, Robert Luallen, Patrick Phillips and members of the Félix lab for their participation in the field. We thank WormBase.

## Author contributions

MAF conceived the project. MAF performed field work and nematode isolation. RT and ECA (CaeNDR) provided genome sequences. XW prepared genomic DNA for sequencing, with assistance from AR. XW and MAF performed the genomic analyses. MAF supervised the project.

