## Supplement for "Mutational divergence over years in local populations of the selfing nematode *Caenorhabditis elegans*"

### Supplementary Materials

**Supplementary Tables**

Tables S1-S6, S9-S11 are separate sheets in the Supplementary Table. (Excel format)
Tables S7 and S8 are below.
File S1 is the code for SNP variant calling.

**Table S1** List of strains with whole-genome sequencing used in this work.
'Strain': the name of strain. 'HS isotype': isotype group (HS for Haplotype Santeuil). 'CaeNDR isotype reference': isotype group in the CaeNDR.org database. 'Category': local, external, or recombinant for HS2 and HS3. 'Year': sampling year. 'Date': sampling date. 'Sample_number.Isolated_nematode': a sample identifier followed by identifier for the isolated nematode from which the strain was derived. 'Location': sampling location (with geographic coordinates). UTM_X: real longitude of each strain in Universal Transverse Mercator (UTM) system (WGS 1984 UTM Zone 31N). UTM_Y: real latitude of each strain in UTM system. UTM_X_projected: the longitude in UTM system of each strain projected onto the simplified straight line. UTM_Y_projected: the latitude in UTM system of each strain projected onto the simplified straight line. One-dimensional Coordinate: the projected location of each strain onto a simplified linear axis (with the strain from the northernmost site designated as 0; values increase southward). UTM coordinates were only shown in local HS2 and HS3 strains. 'Substrate_original': the original substrate information recorded in the field. 'Sample_category': rotting plant, animal, or other. 'Sample_subcategory': detailed description of the substrates.

**Table S2** Isotype proportions in the 2009-2022 Santeuil wood collections.

**Table S3** List of nuclear base substitution mutations identified in HS2 (Santeuil wood set).
Classification: 'external' are SNPs which polymorphism only occurs in external strains; 'recombinant': SNPs which polymorphism only occurs in recombinants; 'local' are SNPs with polymorphism occurring in non-recombinant external strains.

**Table S4** List of nuclear base substitution mutations identified in HS3 (Santeuil wood set).
Classification: external are SNPs with polymorphism occurring only when considering the isotype set including external strains; local are SNPs with polymorphism occurring in the set of local non-recombinant strains.

**Table S5** Distribution of mutation densities among chromosomes, recombination domains and genomic regions for the Santeuil wood set of HS2 (left) and HS3 (right) strains.
Dinucleotide variants were lumped as a single site when calculating SNP density. The table also indicates the number of base pairs and GC content of each category. 'num_snp': number of SNPs in each category. 'len_trimmed': the size of the trimmed genomic region in each category (bp). 'density': the density of SNPs in each category. 'GC content (%)': the proportion of GC of the trimmed genomic region in each category. Below are indicated counts per recombination domain and genomic region for each chromosome.

**Table S6** The counts of each of the six categories of mutations in different genomic regions and mutational bias towards AT for the Santeuil wood set of HS2 and HS3.

**Table S9** Euclidean distance (unit: meter) between strains within HS2. Strains collected from animal carriers are highlighted in bold red text.

**Table S10** Euclidean distance (unit: meter) between strains within HS3. Strains collected from animal carriers are highlighted in bold red text.

**Table S11** Partial discordance of the *C. elegans* genotypes on decomposing stems and on various invertebrates collected on the same stem.
Column labels are as in Table S1, with 'Within-isotype subclade' being the subclade number as shown in Figure 2. D: discordance. C: concordance.

**Table S7** Observed numbers of non-synonymous (N) and synonymous (S) mutations, and the ratio between non-synonymous and synonymous mutations ($R_{N/S}$) in wild isotypes (i.e., HS2 and HS3; highlighted by Wild), and mutation accumulation lines of *Caenorhabditis elegans* cultured in lab (highlighted by MA) from [1](Denver *et al.*, 2012), [2](Denver *et al.*, 2009), and [3](Konrad *et al.*, 2019). None of the comparisons show a significant difference (Chi-Square Tests, $p > 0.05$).

| | Wild HS2 | Wild HS3 | MA PB306 [2] | MA N2 [1] | MA N2 [2] | MA N2 [3] |
|---|---|---|---|---|---|---|
| Observed number of non-synonymous mutation (N) | 90 | 229 | 34 | 56 | 28 | 190 |
| Observed number of synonymous mutation (S) | 39 | 76 | 16 | 24 | 13 | 54 |
| $R_{N/S}$ | 2.31 | 3.01 | 2.12 | 2.33 | 2.15 | 3.52 |

**Table S8** Pairwise Wilcoxon Rank Sum Tests with Bonferroni correction for testing spatial segregation between subclades of HS2 and HS3.

| | HS3.1 | HS3.2 | HS3.3 | HS3.4.1 | HS3.4.2 |
|---|---|---|---|---|---|
| HS3.1 (n = 27) | - | | | | |
| HS3.2 (n = 17) | **0.001** | - | | | |
| HS3.3 (n = 16) | **0.000** | **0.007** | - | | |
| HS3.4.1 (n = 13) | 1.000 | **0.011** | **0.000** | - | |
| HS3.4.2 (n = 10) | 0.053 | 1.000 | **0.030** | 0.192 | - |
| HS3.5 (n = 5) | **0.006** | **0.034** | 1.000 | **0.021** | **0.010** |

n - number of strains used in this test. The test was performed on line coordinates. A single strain was kept per sample, and the four strains at the basal position of HS3.4 (Figure 2) were not included in this test. *P* value less than 0.05 were highlighted in bold.

### Supplementary Figures

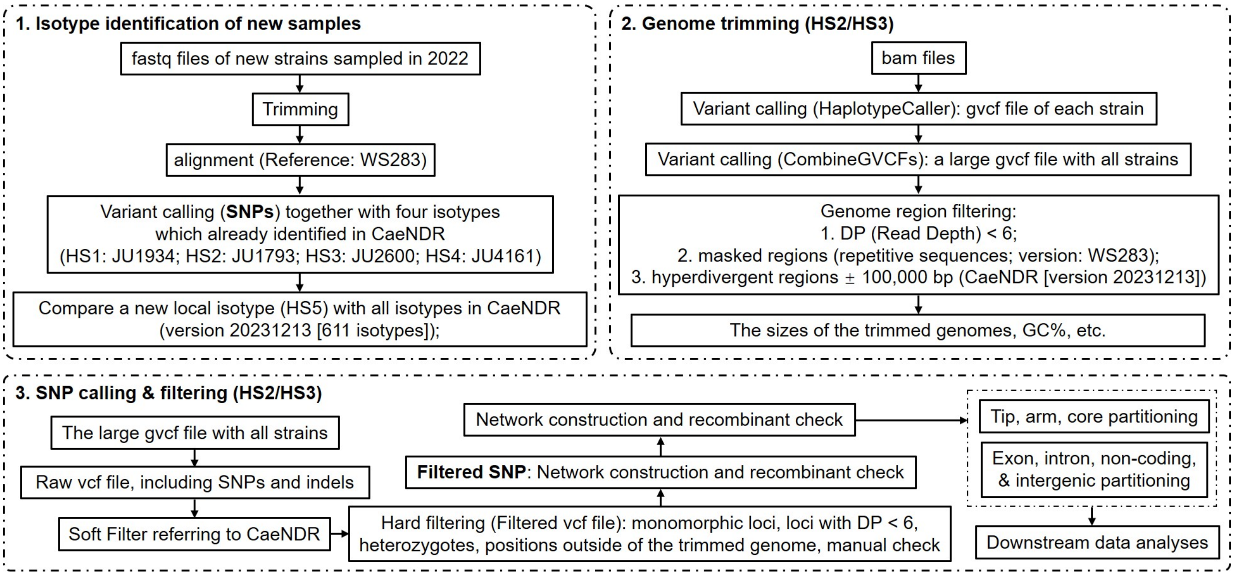


**Figure S1** Flowcharts for isotype identification of strains, genome trimming, and SNP calling and filtering. Genome trimming and SNP calling and filtering were conducted separately in HS2 and HS3.

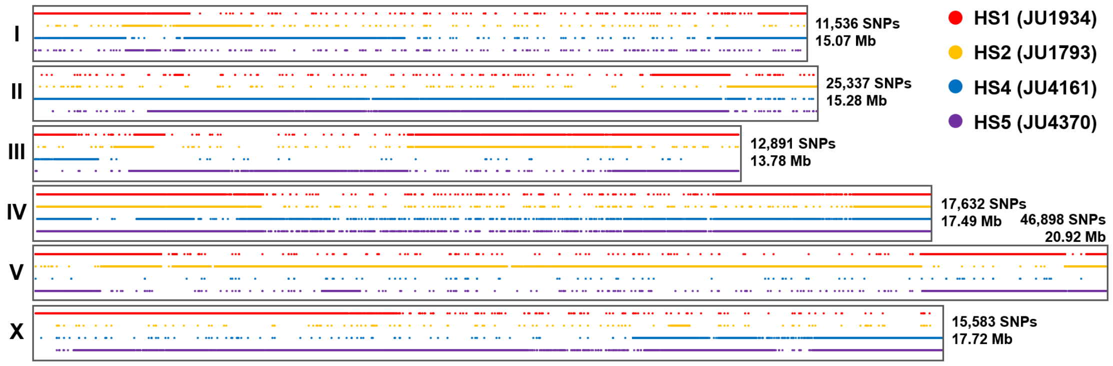


**Figure S2** SNP distribution of representative strains of HS1, HS2, HS4, and HS5 relative to HS3 (JU2600). A continuous line corresponds to numerous adjacent polymorphisms and indicates a high level of divergence with HS3. The genotypes that appeared late in our survey (HS4 and HS5) are not simple recombinants of HS1-3. For example, compared to HS1-3, HS4 carries a divergent center-right region on chromosome I, HS5 on chromosome III or the middle of chromosome X.

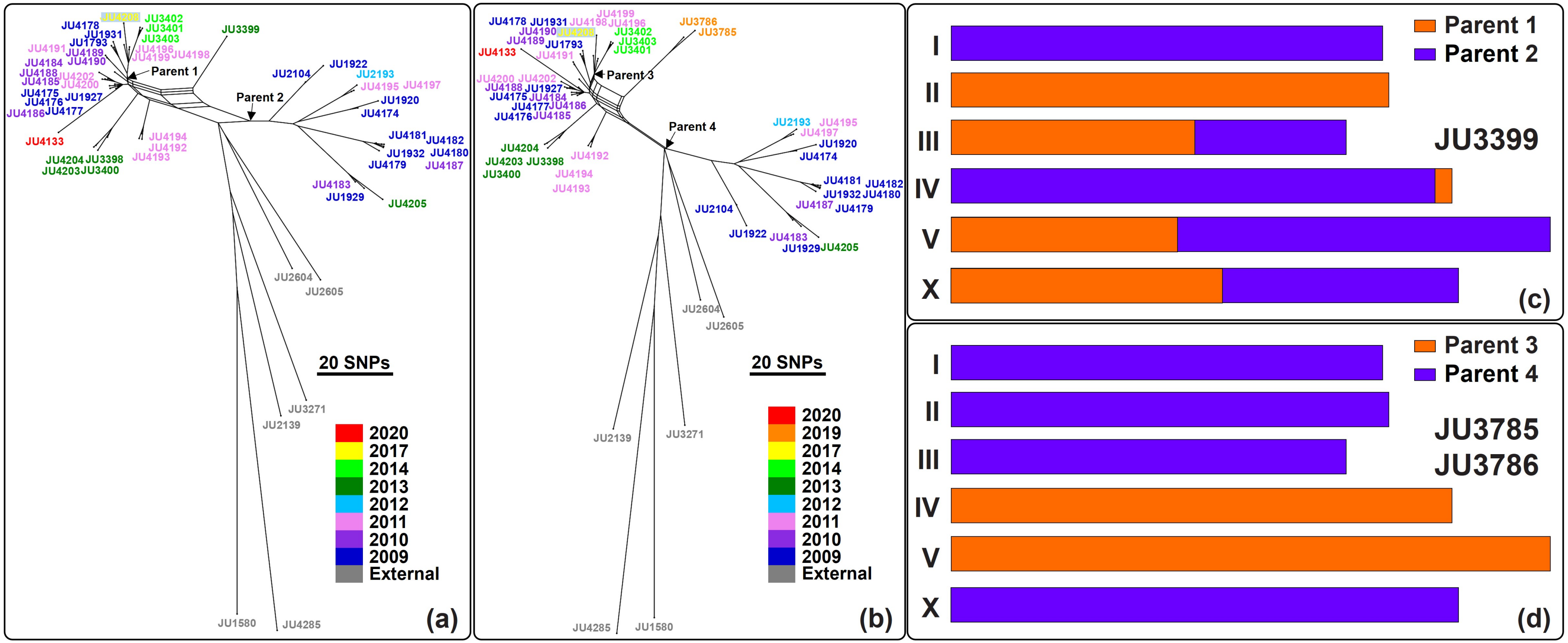


**Figure S3** Recombination within the HS2 genotype. (a) Haplotype network of HS2 with the recombinant JU3399, highlighting the predicted parents of the cross yielding JU3399. (b) Haplotype network of HS2 with the JU3785 and JU3786 recombinants, highlighting the predicted parents of the cross yielding them. (c) Detected chromosomal recombination pattern in the genome of JU3399. (d) Detected chromosomal reassortment pattern in the genomes of JU3785 and JU3786.

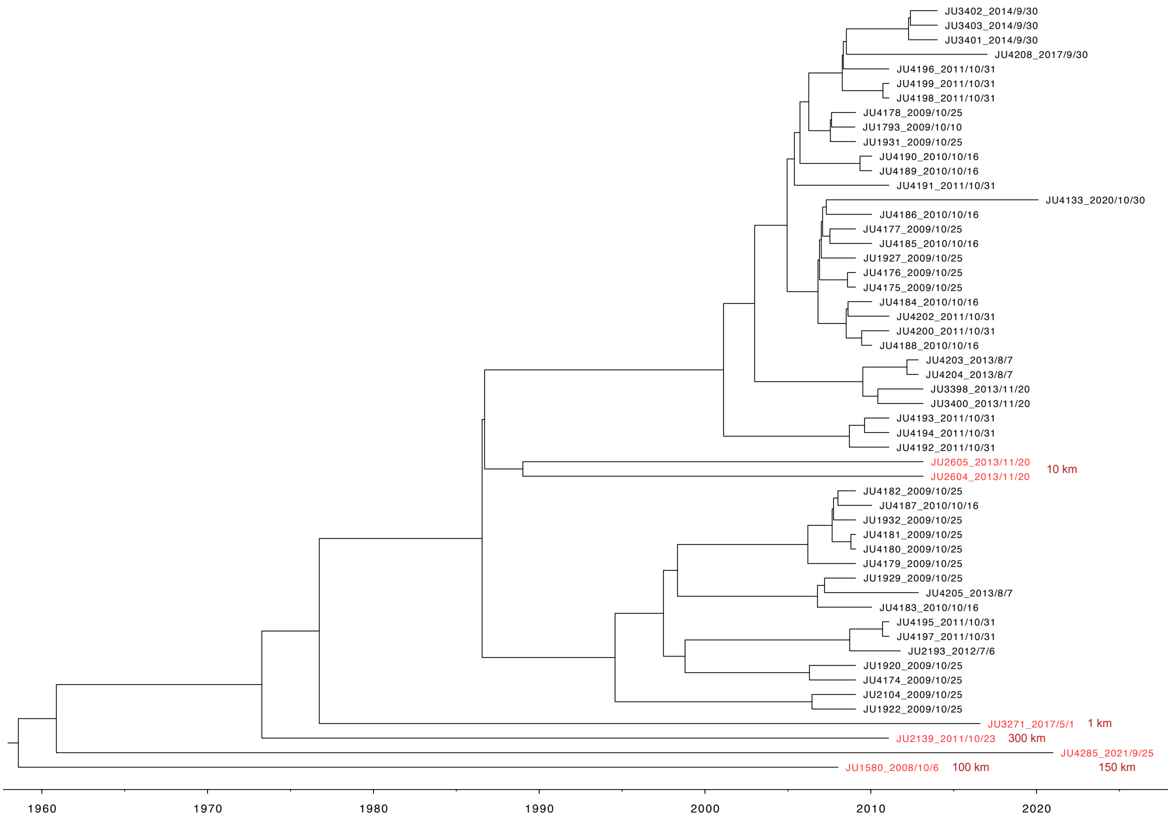


**Figure S4** Maximum Clade Credibility trees of Bayesian Skygrid demographic model with tip-calibration, for the GS2 isotype including external strains from outside the Santeuil wood. Dates of sampling are indicated after the strain name. Approximate distances to the Santeuil site is indicated next to the names of the external strains (in red).

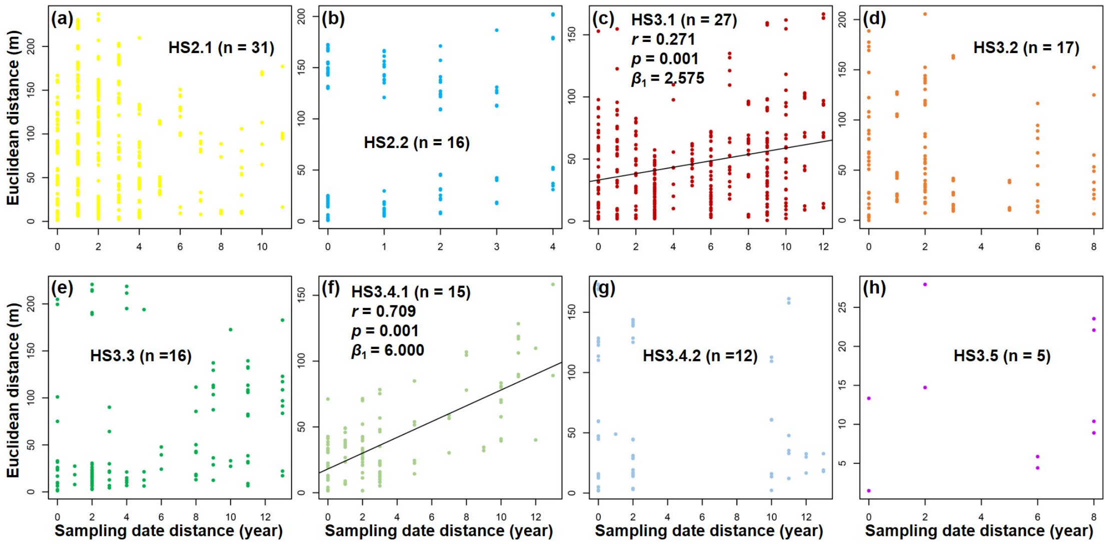


**Figure S5** Euclidean distance versus sampling date distance between strains within each subclade, with significant subclades showing highlighted regression lines, significance (i.e., $p$ value of the Mantel test), $r$ value, and the slope ($\beta_1$). This regression captures directional migration along the transect.

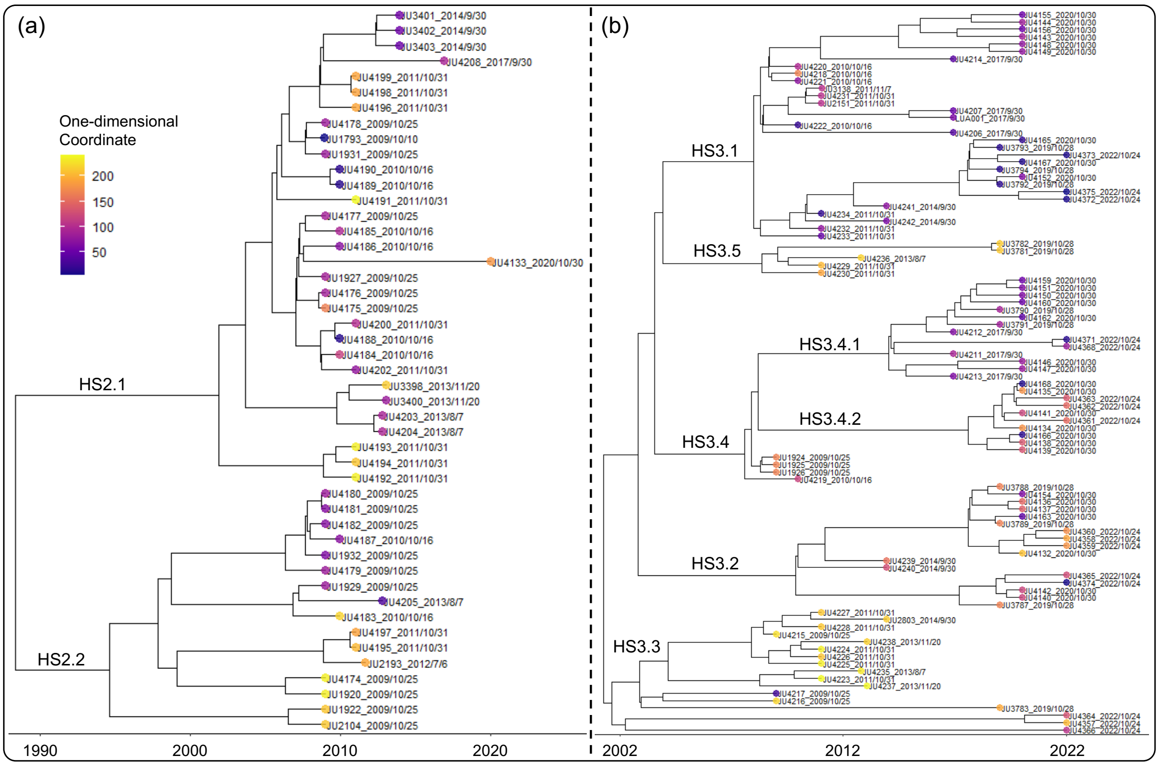


**Figure S6** Spatial positions on the different lineages plotted on the Maximum Clade Credibility trees of Bayesian Skygrid demographic model with tip-calibration for HS2 (a) and HS3 (b). Tip color represents the projected location of each strain onto a simplified linear axis (with the strain from the northernmost site designated as 0 in blue; values in meters increase southward). This figure illustrates the distribution and clustering of strains within each subclade. In some cases, more than one sequenced strain was from the same sample, most notably JU1924-JU1926 in the same stem (Table S1, here at the base of HS3.4). The stem-carrier pairs yielded mostly discordant genotypes (Table S11).
