## Supplementary material for "Mutational divergence over years in local populations of the selfing nematode *Caenorhabditis elegans*": Code: 1_File S1_variant_calling copy.sh

```
#!/bin/bash

source /usr/local/conda/etc/profile.d/conda.sh
conda activate gatk-4.2.2.0

export PATH="$PATH:/users/den/xuwei/bin.linux/nim-2.0.2/bin"
export PATH="$PATH:/users/den/xuwei/bin.linux"

cpus=$1

params_min_depth=5
params_qual=30.0
params_strand_odds_ratio=5.0
params_quality_by_depth=20.0
params_fisherstrand=100.0

params_high_missing=0.95
params_high_heterozygosity=0.10

=/1_data/0_c_elegans_PRJNA13758_WS283/c_elegans.PRJNA13758.WS283.genomic_masked.fa
gvcf=/3_dataset_for_analyses/2_HS2_56_samples/2_2_HS2_get_accurate_region_56_samples/HS2.chr.dp.6.g.vcf.gz
trimmed_region=/3_dataset_for_analyses/2_HS2_56_samples/2_2_HS2_get_accurate_region_56_samples/HS2_no_hyper_regions.merged.bed
genotyping=/3_dataset_for_analyses/2_HS2_56_samples/2_3_HS2_variant_calling_56_samples


bcftools index --tbi ${gvcf}

gatk --java-options "-Xmx40g -Xms1g -XX:ConcGCThreads=${cpus}" \
	GenotypeGVCFs \
	-R ${reference} \
	-V ${gvcf} \
	-O ${genotyping}/HS2.chr.dp.6.vcf.gz

conda activate vcflib_xuweishch

bcftools view -O z ${genotyping}/HS2.chr.dp.6.vcf.gz | het_polarization > ${genotyping}/HS2.chr.bcf
bcftools index ${genotyping}/HS2.chr.bcf
bcftools index --tbi ${genotyping}/HS2.chr.bcf

bcftools view -O v --min-af 0.000001 --threads=${cpus} ${genotyping}/HS2.chr.bcf | \
	vcffixup - | \
	bcftools view -O z -o ${genotyping}/HS2.chr.fixup.vcf.gz
bcftools index ${genotyping}/HS2.chr.fixup.vcf.gz
bcftools index --tbi ${genotyping}/HS2.chr.fixup.vcf.gz

conda activate gatk-4.2.2.0

# soft filter
gatk --java-options "-Xmx4g -Xms1g" \
	VariantFiltration \
	-R ${reference} \
	--variant ${genotyping}/HS2.chr.fixup.vcf.gz \
	--genotype-filter-expression "DP < ${params_min_depth}" --genotype-filter-name "DP_min_depth" \
	--filter-expression "QUAL < ${params_qual}" --filter-name "QUAL_quality" \
	--filter-expression "FS > ${params_fisherstrand}" --filter-name "FS_fisherstrand" \
	--filter-expression "QD < ${params_quality_by_depth}" --filter-name "QD_quality_by_depth" \
	--filter-expression "SOR > ${params_strand_odds_ratio}" --filter-name "SOR_strand_odds_ratio" \
	--genotype-filter-expression "isHet == 1" --genotype-filter-name "is_het" \
	-O ${genotyping}/HS2.chr.soft.filter.1.vcf
bgzip ${genotyping}/HS2.chr.soft.filter.1.vcf
bcftools index --tbi ${genotyping}/HS2.chr.soft.filter.1.vcf.gz

# Apply high missing and high heterozygosity filters
bcftools filter --threads ${cpus} --soft-filter='high_missing' --mode + --include 'F_MISSING <= 0.95' ${genotyping}/HS2.chr.soft.filter.1.vcf.gz | bcftools filter --threads ${cpus} --soft-filter='high_heterozygosity' --mode + --include '( COUNT(GT="het") / N_SAMPLES ) <= 0.10' -O z > ${genotyping}/HS2.chr.soft.filter.vcf.gz

bcftools index ${genotyping}/HS2.chr.soft.filter.vcf.gz
bcftools index --tbi ${genotyping}/HS2.chr.soft.filter.vcf.gz
bcftools stats --threads ${cpus} -s- --verbose ${genotyping}/HS2.chr.soft.filter.vcf.gz > ${genotyping}/HS2.chr.soft.filter.stats.txt

conda deactivate

# Filter loci by bed file
bedtools intersect -a ${genotyping}/HS2.chr.soft.filter.vcf.gz -b ${trimmed_region} -header > ${genotyping}/HS2.chr.soft.filter.v2.merged.vcf.gz

# Filter loci with missing data (for checking)
bcftools view -e 'GT="./."' ${genotyping}/HS2.chr.soft.filter.v2.merged.vcf.gz -O z -o ${genotyping}/HS2_chr_no_missing_data.merged.vcf.gz

# Filter monomorphic loci
bcftools view -i 'GT="1/1"' ${genotyping}/HS2_chr_no_missing_data.merged.vcf.gz | bcftools view -i 'GT="0/0"' -O z -o ${genotyping}/HS2_chr_no_mono.merged.vcf.gz

# Filter heterozygotes (for controlling new mutation in the lab)
bcftools view -g '^het' ${genotyping}/HS2_chr_no_mono.merged.vcf.gz -O z -o ${genotyping}/HS2_chr_no_heterozygotes.merged.vcf.gz

# Filter FORMAT/AD[*:0] = 0 & FORMAT/AD[*:1] = 0 >>> retained
# Filter FORMAT/GT = "0/0" & FORMAT/AD[*:1] != 0 >>> discarded
# Filter FORMAT/GT = "1/1" & FORMAT/AD[*:0] != 0 >>> discarded
# bcftools view -e 'FORMAT/AD[*:0] = 0 & FORMAT/AD[*:1] = 0' ${genotyping}/HS3_chr_no_heterozygotes.vcf.gz | bcftools view -e 'FORMAT/GT = "0/0" & FORMAT/AD[*:1] != 0' | bcftools view -e 'FORMAT/GT = "1/1" & FORMAT/AD[*:0] != 0' -O z -o ${genotyping}/HS3_chr_check.vcf.gz
bcftools view -e 'FORMAT/AD[*:0] = 0 & FORMAT/AD[*:1] = 0' ${genotyping}/HS2_chr_no_heterozygotes.merged.vcf.gz -O z -o ${genotyping}/HS2_chr_check.merged.vcf.gz

# Filter out positions within 50 bp of indels >>> discarded
# bcftools filter -G 50 -g 50 ${genotyping}/HS3_chr_check.vcf.gz -O z -o ${genotyping}/HS3_chr_indel_filter.vcf.gz

# Extract hyperdivergent regions of each isotype from the bed file
# Removes 1 kb on either side of the Cendr hyperdivergent regions
# Remove hyperdivergent regions from vcf file
#grep JU2600 ./20220216_c_elegans_divergent_regions_strain.bed > 6_hyperdivergent_HS3_JU2600.bed # only need to be done one time
#awk '{$2-=100000}1' 6_hyperdivergent_HS3_JU2600.bed | awk '{$3+=100000}1' > 6_hyperdivergent_HS3_JU2600.entended.bed # only need to be done one time
#sed -i 's/ /\t/g' 6_hyperdivergent_HS3_JU2600.entended.bed # only need to be done one time
# for checking
# bedtools intersect -a ${genotyping}/HS3_chr_indel_filter.vcf.gz -b ./hyperdivergent_HS3_JU2600.entended.bed -v -header > ${genotyping}/HS3_chr_no_hyper.vcf.gz
bedtools intersect -a ${genotyping}/HS2_chr_check.merged.vcf.gz -b ${genotyping}/hyperdivergent_HS2_JU1793.extended.20231213.bed -v -header > ${genotyping}/HS2_chr_no_hyper.merged.vcf
bgzip ${genotyping}/HS2_chr_no_hyper.merged.vcf

# Remove duplicates
bcftools norm -d none ${genotyping}/HS2_chr_no_hyper.merged.vcf.gz -O z -o ${genotyping}/HS2_chr_rm_dup.merged.vcf.gz

# Filter FORMAT/DP < 6; filter FORMAT/AD[*:0] + FORMAT/AD[*:1] < 6 (for checking)
bcftools view -e 'FORMAT/DP[*] < 6' ${genotyping}/HS2_chr_rm_dup.merged.vcf.gz | bcftools view -e 'FORMAT/AD[*:0] + FORMAT/AD[*:1] < 6' -O z -o ${genotyping}/HS2_chr_depth_filter.merged.vcf.gz

# Retain only SNPs
bcftools view -v snps ${genotyping}/HS2_chr_depth_filter.merged.vcf.gz -O z -o ${genotyping}/HS2_chr_only_snp.merged.vcf.gz

# Retain only indels
bcftools view -v indels ${genotyping}/HS2_chr_depth_filter.merged.vcf.gz -O z -o ${genotyping}/HS2_chr_only_indel.merged.vcf.gz
```
